# Transcriptional control of central T cell tolerance by NR4A family nuclear receptors

**DOI:** 10.1101/2024.05.19.594881

**Authors:** Hailyn V. Nielsen, James L. Mueller, Ryosuke Hiwa, Letitia Yang, Irena Proekt, Elze Rackaityte, Dominik Aylard, Christopher D. Scharer, Mark Anderson, Byron Au-Yeung, Julie Zikherman

**Affiliations:** Division of Rheumatology, Rosalind Russell and Ephraim P. Engleman Arthritis Research Center, Department of Medicine, University of California, San Francisco, CA, 94143; Department of Rheumatology and Clinical Immunology, Graduate School of Medicine, Kyoto University, 54 Kawahara-cho, Shogoin, Sakyo-ku, Kyoto 606-8507, Japan; Biomedical Sciences Graduate Program, University of California, San Francisco, CA, 94143; Diabetes Center, Department of Medicine, University of California, San Francisco, CA, 94143; Department of Biochemistry and Biophysics, University of California, San Francisco, CA, 94143; Department of Molecular & Cell Biology, University of California, Berkeley, CA, 94720; Department of Microbiology and Immunology, Emory University, Atlanta, GA 30322, USA; Division of Immunology, Lowance Center for Human Immunology, Department of Medicine, Emory University, Atlanta, GA 30322, USA

**Author notes:** Corresponding authors: Dr. Julie Zikherman 513 Parnassus Avenue, Room HSW1201E, Box 0795 San Francisco, CA 94143-0795 E-mail address; Dr. Byron Au-Yeung Emory University 615 Michael Street, Whitehead Biomedical Research Building, Room 246 Atlanta, GA 30322.

**Keywords:** NUR77, NOR-1, *Nr4a1*, *Nr4a3*, Treg, T cell tolerance, anergy, negative selection, clonal deletion, tissue restricted antigen (TRA), medullary thymic epithelial cell (mTEC), recent thymic emigrant (RTE), *Bcl2l11*/BIM, *Ikzf2*/HELIOS, *Nt5e*/CD73

## Abstract

Although deletion of self-reactive thymocytes and their diversion into regulatory T cell (Treg) lineage are critical for immune tolerance and homeostasis, the molecular pathways that link antigen recognition to these fates are incompletely understood. The Nr4a nuclear hormone receptors are transcriptionally upregulated in response to TCR signaling in the thymus and are implicated in both deletion and diversion, but the mechanisms by which they operate are not clear. Redundancy among the family members and their requirement for Treg generation and maintenance have obscured their role in negative selection. Here we take advantage of competitive bone marrow chimeras and the OT-II/RIPmOVA model to demonstrate that *Nr4a1* and *Nr4a3* are essential for upregulation of *Bcl2l11*/BIM and negative selection by tissue-restricted model self-antigen (TRA). Moreover, we reveal that the Nr4a family is absolutely required for full induction of a broad transcriptional program triggered in self-reactive thymocytes by TRA recognition, and conserved across model systems and the natural repertoire. Importantly, both model self antigen-specific TCR Tg and polyclonal thymocytes lacking *Nr4a1/3* that escape negative selection acquire an anergy-like program that persists in the periphery and is also evident among wild-type recent thymic emigrants (RTEs). We propose that the Nr4a family transduces TCR signals during thymic development to enforce the fates of highly self-reactive clones, mediating not only deletion and Treg diversion, but also contributing to a cell-intrinsic, persistent anergy-like program that may operate at the margins of canonical thymic tolerance mechanisms to restrain self-reactive T cells after thymic egress.

## Introduction

Deletion of highly self-reactive thymocytes and diversion of others into the regulatory T cell lineage during their development are vital to prevent autoimmune disease. Indeed, mice and humans with defects in these central T cell tolerance mechanisms exhibit a broad range of autoimmune pathology^1-3^. Positive selection by specialized cortical thymic epithelial cells (cTECs) ensures MHC restriction among double positive (DP) thymocytes^4^. Although excessive recognition of self-pMHC in the thymic cortex can trigger deletion of DP thymocytes at this early stage, many important self-antigens are tissue-restricted^4-6^. Only after their migration to the thymic medulla following positive selection are thymocytes tolerized to such tissue-restricted antigens^4,5^. Promiscuous gene expression by AIRE-positive medullary thymic epithelial cells (mTECs) along with recently discovered thymic mimetic cells serves to eliminate self-reactivity from the conventional mature T cell repertoire^2,6-8^. In contrast to the thymic cortex, medullary self-antigen encounter can drive both Treg diversion and deletion^5^. However, the molecular pathways that direct these fate decisions are incompletely understood. Although the pro-apoptotic BH3-only family member BIM is implicated in deletion of self-reactive thymocytes^9^, the mechanisms linking it to self-antigen recognition are not clear. In contrast to the discrete fates of death and Treg diversion, TCR affinity for a given self-antigen exists on a spectrum, highlighting the need for a cell-intrinsic restraint on clones that enter the mature conventional repertoire with a TCR affinity that falls just below the thymic signaling threshold for death or diversion.

Among the few factors directly implicated in thymic negative selection are the Nr4a family of orphan nuclear hormone receptors. *Nr4a1, Nr4a2, and Nr4a3* (encoding Nur77, Nurr, and Nor1 respectively) comprise a small family of transcriptional regulators with conserved DNA-binding domains and shared binding site sequence, but lacking well-defined ligands. Rather, the Nr4a family member are primary response genes that are rapidly induced by antigen receptor signaling and thought to be regulated at the level of their dynamic expression. They were first identified as mediators of antigen-induced cell death in T cell hybridomas, and studies of full length and truncated dominant negative transgenes have suggested roles in thymic negative selection ^10-14^. However, the relatively subtle phenotypes of *Nr4a1* and *Nr4a3* single knock-out mice raises the possibility that redundancy among Nr4a family members masks their role in thymic selection ^15-18^. Conversely, mice lacking *both Nr4a1* and *Nr4a3* in the germline or T cell lineage exhibit loss of Treg with resultant severe immune dysregulation and death by 4 weeks of age^19-22^. Therefore, dissecting the roles of the Nr4a family during thymic negative selection and dissecting the mechanisms by which they operate has been challenging.

To overcome this challenge, we recently used a competitive chimera strategy to reconstitute a normal Treg compartment of WT genetic origin, and to unmask redundant roles of *Nr4a1* and *Nr4a3* in tolerance ^19^. We found that polyclonal double knockout (DKO) thymocytes lacking both *Nr4a1* and *Nr4a3* exhibit a profound competitive advantage between DP and SP stages of thymic development, suggesting escape from negative selection. However, the anatomical site and developmental stage where Nr4a-dependent deletion occurs, as well as the molecular mechanisms involved, remain unclear. Indeed, both genomic and non-genomic Nr4a-dependent pathways to apoptosis have been proposed, but transcriptional targets that mediate deletion are unknown. Here we take advantage of the OT-II TCR Tg model to track Nr4a-deficient and Nr4a-sufficient thymocytes as they develop in the presence or absence of cognate model self-antigen (chicken ovalbumin) expressed by mTECs (RIPmOVA Tg). We show that Ag-specific DKO thymocytes escape mTEC-dependent negative selection and that these thymocytes fail to upregulate *Bcl2l11* transcript and BIM protein in response to self-antigen recognition. We further show that the Nr4a family is required to impose a broad transcriptional program triggered by self-antigen recognition in semimature thymocytes and conserved across Tg models and in polyclonal settings. DKO thymocytes destined for negative selection instead escape deletion and engage an anergy-associated program that is also evident among naïve peripheral T cells. Importantly, we show that phenotypically anergic DKO SP thymocytes are massively expanded even in polyclonal chimeras with an unrestricted TCR repertoire. Nr4as therefore mediate deletion, Treg diversion, and contribute to a non-deletional tolerance transcriptional program initiated in the thymus that can persist in the periphery and may serve as a fail-safe mechanism to prevent frank autoimmunity.

## Results

### TCR signaling upregulates Nr4a family in thymocytes destined for deletion and Treg diversion

*Nr4a* transcripts are rapidly but transiently induced following Ag stimulation^23,24^. *Nr4a1* transcript is much more abundant than *Nr4a2* and *Nr4a3* in the T cell lineage (**Supp Fig 1a**; Immgen.org). Mass spectrometry confirms that all three family members are rapidly upregulated in mature CD8+ cells by TCR stimulation, yet only Nr4a3 protein persists without decline over 24 hours, implying a longer protein half-life and suggesting levels may accumulate despite low abundance transcript (**Supp Fig 1b**; immpres.co.uk). Conversely, although Nr4a1/Nur77 protein is most highly induced, it is rapidly degraded ^25-27^.

Previously described reporters of *Nr4a* transcript expression, including Nur77/*Nr4a1*-GFP BAC Tg, reveal sensitivity to affinity, dose, and duration of TCR stimulation and upregulation at signal dependent checkpoints during thymic development ^23,24,28-32^. Nur77-GFP is upregulated in post-selection DP thymocytes concurrently with CD69 induction (**Fig 1a, b**), increases at the semimature SP stage^33^, and peaks in agonist-selected Treg cells – consistent with their high degree of self-reactivity (**Fig 1a, Supp Fig 1c, d**).

**Figure 1.**
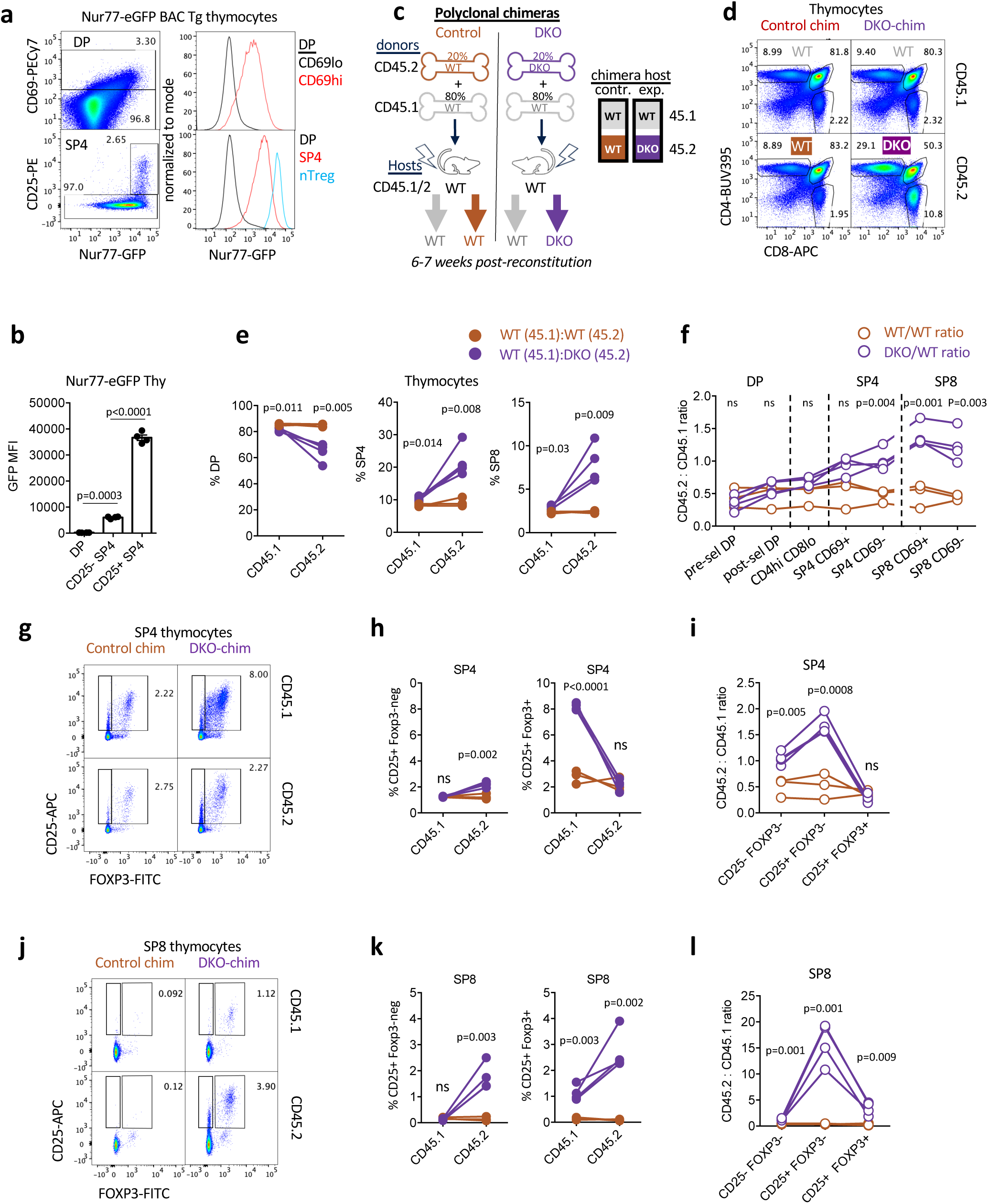
Polyclonal DKO T cells exhibit a competitive advantage during thymic selection. **a, b.** Nur77-eGFP BAC Tg thymocytes were stained to detect thymic subsets. Lefthand panels depict representative gating to identify CD69hi post-selection DP thymocytes and CD25hi SP4 Treg. Righthand histograms depict GFP expression in gated populations. Graph in b depicts GFP MFI quantification of 4 biological replicates +/- SEM. Data **c.** Schematic depicts radiation chimera design. **d.** Thymocytes from chimeras were stained to detect congenic markers CD45.1 and 2 as well as CD4/CD8 co-receptors. Representative plots depict thymic subsets of each donor genotype. **e.** Graphs depict quantification of gating in (d) with lines connecting donor genotypes from individual chimeras. **f.** Graph depicts ratio of CD45.2/CD45.1 donor genotypes from individual chimeras within DP and SP populations further sub-gated to define CD69hi and CD69lo compartments. **g-l.** Representative plots depict SP4 (g) or SP8 (j) thymocytes of each donor genotype from both experimental and control chimeras (as above) stained to identify CD25 and Foxp3 expression. Graphs in h, k depict quantification of gating in g, j respectively with lines connecting donor genotypes from individual chimeras. Graphs in i, l depict ratio of CD45.2/CD45.1 donor genotypes from individual chimeras within same SP4 and SP8 subset gates. <u>statistical tests:</u> b. one-way ANOVA with Tukey’s multiple hypothesis correction. No matching or pairing. Data in a, b representative of at least 4 biological replicates e, h, I, k, l. Unpaired t tests, Do not assume same SD f. Multiple unpaired T tests, Holm-Sidak method for multiple comparison correction without assuming consistent SD. Graphs except b depict N=3 WT chimeras and N=4 DKO chimeras. Data in a are representative of more than 3 independent experiments. Data in d-i representative of at least 3 independent sets of chimeras.

*Nr4a1* and *Nr4a3* transcripts were previously reported to be upregulated in thymocytes undergoing negative selection ^34,35^. Most recently, a CITEseq atlas of thymic development captured two clusters of thymocytes destined for deletion (**Supp Fig 1e**)^36,37^. One of these clusters corresponds to CD69hi DP thymocytes with low *Ccr7* transcript levels and may represent cells undergoing an initial wave of deletion triggered by excessive self-pMHC recognition in the cortex prior to CCR7-dependent migration to the medulla^38-40^. The second cluster is closely apposed to Foxp3-expressing Treg and captures CD69hi semimature SP thymocytes with high *Ccr7* transcript levels, likely those undergoing deletion in response to promiscuous expression of tissue-restricted antigens (TRA) by medullary thymic epithelial cells (mTECs) (**Supp Fig 1e**)^6,41,42^. Notably, *Nr4a1* and *Nr4a3* transcripts are highly correlated with both deleting clusters, while *Nr4a2* is detectable in only a sparse number of cells (**Supp Fig 1e**).

### Nr4a-deficient thymocytes exhibit a competitive advantage during development

We recently developed a competitive radiation chimera strategy in which donor bone marrow (BM) lacking both *Nr4a1* and *Nr4a3* (DKO) is mixed with an excess of WT BM to reconstitute a normal Treg compartment of WT origin (**Fig 1c**) ^19^. Chimeras reconstituted with a congenically marked WT:WT donor BM mix with matched input ratio serve as controls.

DKO cells exhibit a large cell-intrinsic advantage relative to WT among SP4 and SP8 thymocytes in the context of normocellular chimeras (**Fig 1d, e, Supp Fig 2a**). This advantage begins – albeit to a subtle degree – after the positive selection checkpoint at the CD69hiTCRβhi DP stage (**Supp Fig 2b, c**, normalized ratios). This might reflect either enhanced positive selection or impaired negative selection. However, the most profound advantage for DKO thymocytes in competitive chimeras is most apparent at the SP stages of thymic development (**Fig 1e, f, Supp Fig 2d, e)**.

Strikingly, DKO cells exhibited a much larger advantage in the SP8 CD69hi stage than in the SP4 CD69hi stage (**Fig 1f**). This may reflect greater escape of SP8 than SP4 from deletion in the medulla. Indeed, we observe progressively reduced Nur77-GFP reporter expression during normal SP8 maturation compared to SP4 maturation (**Supp Fig 1d**), which could be consistent with more stringent pruning of this compartment^43,44^.

### Nr4as regulate Foxp3+ populations in cell intrinsic and non-cell intrinsic manner

DKO:WT chimeras reconstituted a Treg compartment of largely WT origin - as previously reported (**Fig 1g-i, Supp Fig 2f-h**)^19^. However, the total thymic Treg (tTreg) compartment was expanded in DKO:WT chimeras relative to WT:WT chimeras (**Fig 1g, h, Supp Fig 2f**). This cell non-intrinsic phenotype might be mediated by enhanced IL-2 production by self-reactive DKO thymocytes ^5,45-47^. Indeed, CITEseq data reveals an IL-2/STAT5 signature overlapping with highly signaled SP cluster destined for deletion/diversion (**Supp Fig 1e**). CD25+Foxp3-SP4 population are proposed to represent precursors of cells that can either be deleted or recruited to Foxp3 lineage^5,45^. Although DKO thymocytes were depleted in the SP4+Foxp3+CD25hi gate relative to WT cells from the same chimera, they were dramatically expanded among this CD25hi Foxp3-population (**Fig 1g-i)**).

Unexpectedly, we discovered a novel CD25+SP8 population in DKO:WT experimental chimeras that was undetectable in WT:WT control chimeras and almost exclusively populated by DKO thymocytes (**Fig 1j-l**). We also identified a SP8+Foxp3+CD25hi population only in DKO:WT chimeras and enriched among DKO thymocytes (**Fig 1j-l, Supp Fig 2i-j**). An analogous population of Foxp3+ CD8 T cells have been described among tumor infiltrating lymphocytes and in GVHD. Strikingly, these peripheral cells are expanded in the absence of BIM suggesting this fate may be normally censored by deletion^48,49^.

### *Nr4a1* and *Nr4a3* are required for negative selection by tissue-specific Ag

We next sought to test the hypothesis that Nr4a family members mediate antigen-dependent negative selection of thymocytes in the medulla. To model thymocyte recognition of self-antigen expressed by mTECs, we took advantage of OT-II TCR Tg that recognizes OVA 323-339 peptide presented by Class II MHC. We generated BM chimeras with hosts expressing RIPmOVA Tg in which the rat insulin promoter directs expression of a model self-antigen (membrane-tethered ovalbumin) in pancreatic beta islet cells (and in renal proximal tubular cells and testes), as well as in radioresistant mTECs ^50,51^. In these chimeras, OT-II and DKO OT-II thymocytes could be tracked through development in the presence of cognate antigen expression by mTECs (**Fig 2a**). Importantly, to provide an abundant source of normal Treg, we included congenically-marked polyclonal WT donor BM (**Fig 2a**).

**Figure 2.**
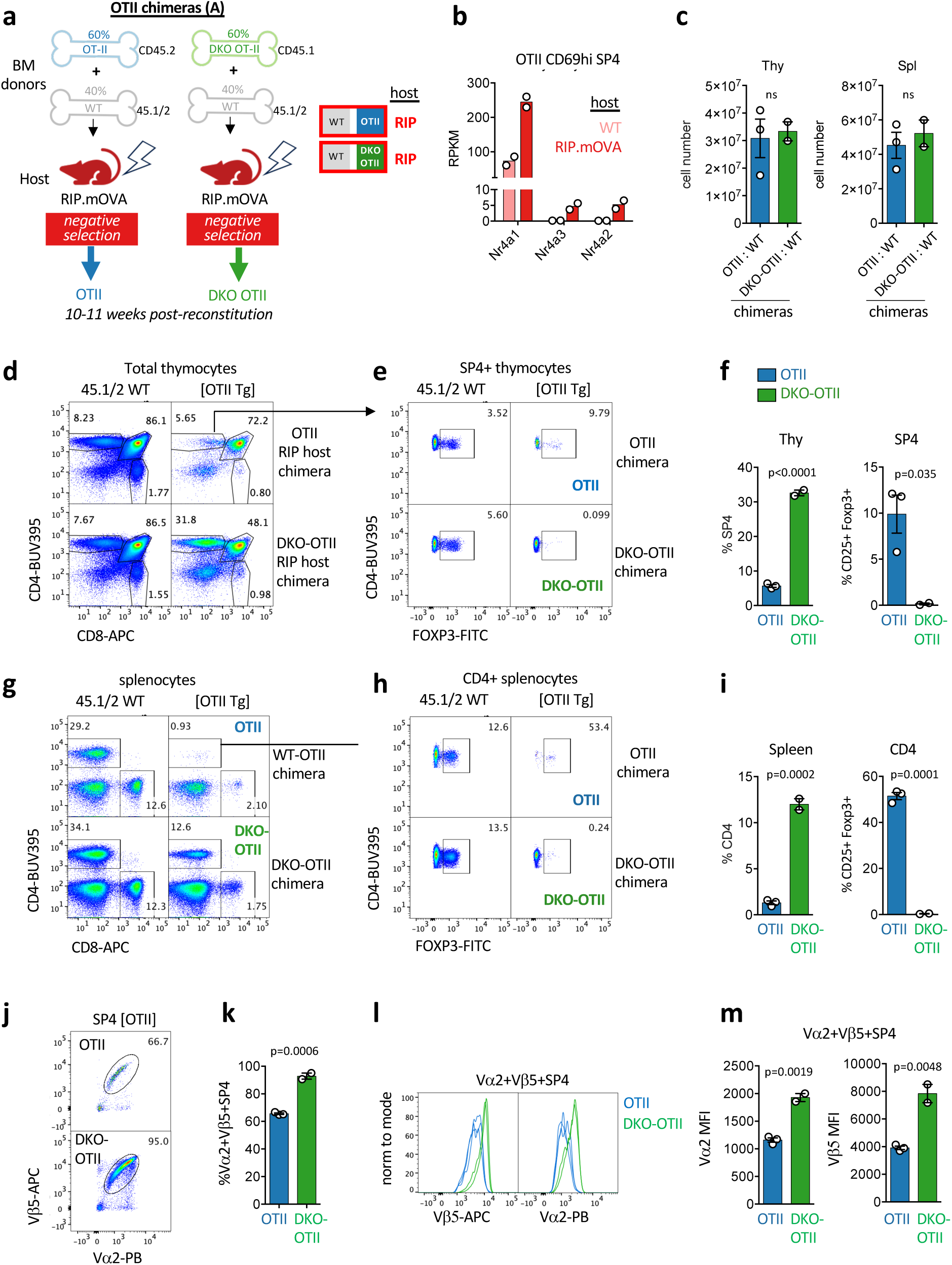
*Nr4a1* and *Nr4a3* are required for negative selection by tissue-specific Ag. a. Schematic depicts radiation chimera design. b. Graph depicts rpkm of *Nr4a1-3* transcripts in CD69hi SP4 va2+vb5+ OTII thymocytes sorted from either WT or RIPmOVA host chimeras in biological replicate (Fig 3a schematic) and subjected to RNAseq (Table 1a depicts p value and fdr for WT vs RIPmOVA hosts, GSE235101). c. Quantification of total thymus and spleen cellularity from chimeras depicted in (a) were harvested 10-11 weeks post irradiation and BM transfer. Values are plotted from individual chimeras +/- SEM **d-i.** thymocytes (d-f) and splenocytes (g-i) from chimeras depicted in (a), harvested as in (c), and stained to detect congenic markers CD45.1/2, CD4/8 co-receptors, and intra-cellular Foxp3 expression. Representative plots depict thymic or peripheral splenic T cell subsets of each donor genotype (d, g), Treg gating among SP4/CD4 T cells of each donor genotype (e, h), and quantification of these values from N=2 DKO-OTII/RIPmOVA and N=3 WT-OTII/RIPmOVA chimeras (f, i). **j-m.** Thymocytes from these RIPmOVA chimeras (a) were stained to detect vβ5 and vα2 TCR surface expression among subsets. Representative plots depict vβ5+ vα2+ OTII TCR gate among SP4 thymocytes of either WT-OTII or DKO-OTII donor genotype, and quantification of these values +/- SEM (j, k). Representative histograms depict surface expression of these TCR chains from SP4 thymocytes of either donor genotype and quantification of MFI +/- SEM (l, m). <u>Statistical tests:</u> c, unpaired two-tailed parametric t-tests f, i, k, m, unpaired two-tailed non-parametric t-tests Graphs depict N=2 DKO-OTII/RIPmOVA and N=3 WT-OTII/RIPmOVA chimeras and are representative of at least 3 biological replicates of each. Data in d-m are representative of 3 independent sets of OTII/RIPmOVA chimeras.

We found that semimature CD69hi SP4 OT-II (Vα2+Vβ5+) thymocytes expressed only *Nr4a1* in WT hosts lacking model self-antigen, while all three *Nr4a* transcripts were induced in negatively-selecting RIPmOVA hosts, although absolute abundance of *Nr4a1* far exceeded that of *Nr4a2* and *Nr4a3* (**Fig 2b**). At 10-11 weeks post-reconstitution, nearly all OT-II SP4 thymocytes in RIPmOVA hosts are efficiently deleted^51,52^ (**Fig 2d, e, f**); very few OT-II cell remain, some of which are diverted towards the thymic Treg compartment (CD25hi Foxp3-precursors and CD25hi Foxp3+ Treg) (**Fig 2e, f, Supp Fig 3a-c**). Similarly, OT-II CD4 T cells are largely absent from the periphery of RIPmOVA chimeras with half of the few remaining cells expressing Foxp3 (**Fig 2g, h, i, Supp Fig 3d**). By contrast, DKO OT-II SP4 thymocytes in RIPmOVA hosts evade both deletion and Treg diversion and readily populate the periphery (**Fig 2d-i, Supp Fig 3a-d**).

Counterselection of Vα2+Vβ5+ OT-II thymocytes in RIPmOVA hosts is accompanied by downregulation of surface TCR expression (**Fig 2j-m**). DKO OT-II escape deletion and evade downregulation of surface TCR Vα2 and Vβ5 chains even in the face of RIPmOVA cognate self-antigen expression in the thymus (**Fig 2j-m**). Together, these data suggest that *Nr4a1* and *Nr4a3* are absolutely required for negative selection of thymocytes in the OT-II / RIPmOVA model of deletion by TRA.

### *Nr4a1* and *Nr4a3* mediate negative selection of CD69hi SP thymocytes

RIPmOVA host chimeras assessed at relatively late time points following reconstitution exhibited near-complete deletion of OT-II thymocytes, consistent with prior observations that this TCR Tg model favors deletion rather than Treg induction^51^. To assess cell-intrinsic and antigen-dependent effects of the Nr4a family *before* complete deletion of OT-II cells, we analyzed a different set of competitive chimeras at an earlier timepoint. Into either WT or RIPmOVA hosts, we introduced OT-II and DKO OT-II donor BM admixed with an excess of polyclonal congenically marked donor marrow (to reconstitute a Treg compartment of WT origin, as before) (**Fig 3a**). We limited precursor frequency of OT-II thymocytes to avoid overwhelming tolerance machinery^53^. Importantly, we analyzed thymic development 5-7 weeks post-irradiation, prior to complete deletion of virtually all OT-II cells in RIPmOVA hosts.

**Figure 3.**
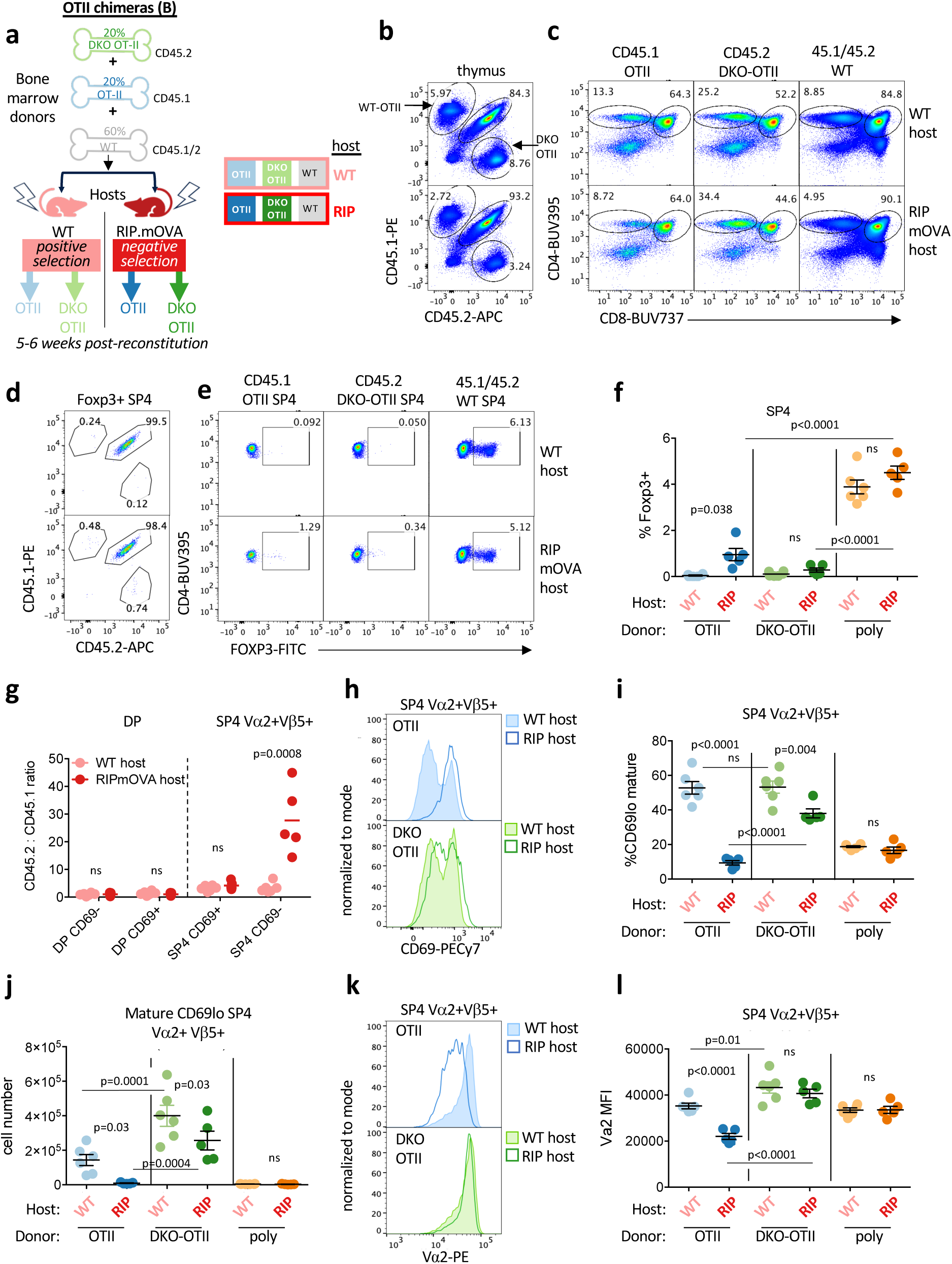
*Nr4a1* and *Nr4a3* mediate negative selection of CD69hi SP thymocytes. **a.** Schematic depicts radiation chimera design. **b, c.** representative plots depict thymocytes from chimeras in (a) harvested 5-6 weeks post irradiation and BM transfer and stained to detect (b) donor genotype and (c) thymic subsets among each donor genotype from WT host (top) or RIPmOVA host (bottom) Statistical tests: f, i, l, one way ANOVA with Tukey’s multiple hypothesis test. g, unpaired two-tailed t-test, do not assume equal SD, corrected for multiple comparisons with Holm-Sidak method. **d-f.** thymocytes from chimeras harvested as above were stained to detect intracellular Foxp3 expression in addition to congenic markers and co-receptors. representative plots depict (d) Foxp3+ SP4 thymocytes stained to detect donor genotype and (e) SP4 thymocytes of each donor genotype (as gated in c) stained to detect Foxp3+ cells. Graph (f) depicts quantification of gates in (e) +/- SEM. **g.** Graph depicts ratio +/- SEM of CD45.2/CD45.1 donor genotypes from individual chimeras within DP and SP populations further sub-gated to define CD69hi and CD69lo compartments. **h-j.** Representative overlaid histograms depict CD69 expression among SP4 Vβ5+ Vα2+ thymocytes of each OTII donor genotype from N=3 WT hosts and N=3 RIPmOVA hosts as color-coded in (a). % (i) and absolute number (j) of CD69lo cells (based on bimodal CD69 distribution in top panel of h) are quantified +/- SEM. k, l. Representative overlaid histograms depict surface expression of TCR Vα2 among Vβ5+ Vα2+ SP4 thymocytes of each OTII donor genotype from N=3 WT hosts and N=3 RIPmOVA hosts as color-coded in (a). Graph depicts quantification of Vα2 MFI in this population +/- SEM. <u>Statistical tests:</u> f, i, j, l one way ANOVA with Tukey’s multiple hypothesis test. g, multiple unpaired t-tests with Holm-Sidak correction for multiple comparisons, do not assume consistent SD across comparisons. Graphs in this figure depict N=6 WT host and N= 5 RIPmOVA host chimeras. Data in this figure are representative of 2 independent sets of chimeras.

In both WT and RIPmOVA hosts, a small and comparable fraction of OT-II and DKO OT-II cells populate a normocellular thymus relative to polyclonal donor cells (**Figure 3b**, **Supp Fig 4a, b**). OT-II thymocytes of either genotype represent a small proportion of the DP gate, yet they outcompete polyclonal thymocytes into the SP4 compartment of WT hosts (**Fig 3c**, **Supp Fig 2c, d, e**). This may reflect more efficient positive selection of OT-II and/or less negative selection relative to polyclonal cells. DKO OT-II cells exhibit a marked competitive advantage into the SP4 compartment relative to polyclonal cells irrespective of RIPmOVA expression (**Fig 3c, Supp Fig 4c-f**). By contrast OT-II thymocytes are modestly reduced in the SP4 compartment in RIPmOVA hosts relative to WT hosts, yet they are not deleted entirely as in Fig 2, leaving them amenable to further study (**Fig 3c, Supp Fig 4c-f)**.

Importantly, only a very small fraction of OT-II is driven into the Treg compartment at this time point (**Fig 3d-f, Supp Fig 4c**). Instead, Treg in this model predominantly originate from polyclonal WT donor BM in either WT or RIPmOVA hosts, both of which have comparable percentage and absolute number of thymic Treg (**Fig 3d-f, Supp Fig 4b, c**). This enabled us to eliminate Treg diversion as a fate and study thymocytes destined for deletion in RIPmOVA hosts.

We next sought to define the stage at which OT-II and DKO OT-II development diverge in these chimeras. CD69 is upregulated at the positive selection checkpoint in DP thymocytes, remains high on semi-mature SP thymocytes and is then downregulated in mature SP thymocytes prior to thymic export (**Supp Fig 1e**)^54^. In RIPmOVA hosts, counter-selection of OT-II thymocytes occurs between the SP4 CD69hi and the mature SP4 CD69lo stage (**Figure 3g**). This is consistent with CITEseq data showing that CCR7hi SP cluster destined for negative selection in the medulla express high levels of CD69 protein on their surface (**Supp Fig 1e**). Indeed, almost no surviving OT-II mature CD69lo SP4 thymocytes can be retrieved from RIPmOVA hosts (**Figure 3h-j**). By contrast, DKO OT-II thymocytes evade counter-selection in RIPmOVA hosts and populate the mature CD69lo SP4 compartment (**Fig 3h-j**). Importantly, the periphery reveals abundant DKO OT-II but near-complete loss of OT-II CD4 T cells in RIPmOVA hosts – as described later in this report (see **Fig 9**). Unexpectedly, DKO OT-II thymocytes exhibit modest advantage relative to OT-II even in WT hosts which may reflect differential response to endogenous pMHC (**Fig 3c, j, Supp Fig 4c, e, f**). Importantly, this advantage is dramatically enhanced in RIPmOVA hosts, confirming dependence of the phenotype on mTEC expression of antigen (**Fig 3i, j**).

Finally, even at this early time point, OT-II SP thymocytes in RIPmOVA hosts already exhibit marked downregulation of surface Vα2+Vβ5+ TCR levels, while DKO OT-II thymocytes retain high surface TCR expression (**Fig 3k, l, Supp Fig 4g, h**) This model enabled us to capture CD69hi SP4 OT-II cells before they are lost to deletion and without diversion towards Treg fate, and therefore served as a unique platform to define mechanism by which Nr4as regulate thymic deletion in the medulla via transcriptional profiling.

### *Nr4a1* and *Nr4a3* are required for thymic transcriptional upregulation of *Bcl2l11*/BIM by self-antigen recognition

The pro-apoptotic BH3-only domain Bcl2 family member BIM is among the few proteins with a well-defined functional role during thymic negative selection ^9,55^. BIM is transcriptionally upregulated by TCR signaling and operates by triggering mitochondrial cytochrome c release and caspase activation^55,56^. T cells from *Bcl2l11/*BIM-deficient mice escape from negative selection by superantigen, peptide injection, and in the HY model of ubiquitous antigen expression ^9,18^. In addition, BIM-deficient SP4 thymocytes escape deletion in the InsHEL model in which HEL antigen is expressed in mTECs, similar to RIPmOVA^37,57^. CITEseq data revealed that *Bcl2l11* transcript is highly expressed among thymocytes undergoing negative selection at the signaled DP stage in the cortex (CCR7-negative) and subsequently in CCR7+ CD69hi semimature SP thymocytes in the medulla, and correlated with expression patterns of *Nr4a1* and *Nr4a3* (**Fig 4a, Supp Fig 1e**). We found that BIM protein expression is upregulated in post-selection DP and in SP thymocytes in proportion to Nur77-GFP reporter expression (**Fig 4b, c, Supp Fig 5a**). Consistent with prior reports, post-selection DP and SP thymocytes are expanded in the absence of BIM, and Nur77-eGFP reporter expression among those populations is markedly increased - consistent with rescue of strongly signaled thymocytes from deletion (**Fig 4d, e**)^30,58^. Taken together, these results suggest that the highest Nr4a1-expressing thymocytes express BIM levels high enough to bind and sequester anti-apoptotic Bcl2 family members in those cells, thereby tipping the balance in favor of apoptosis.

**Figure 4.**
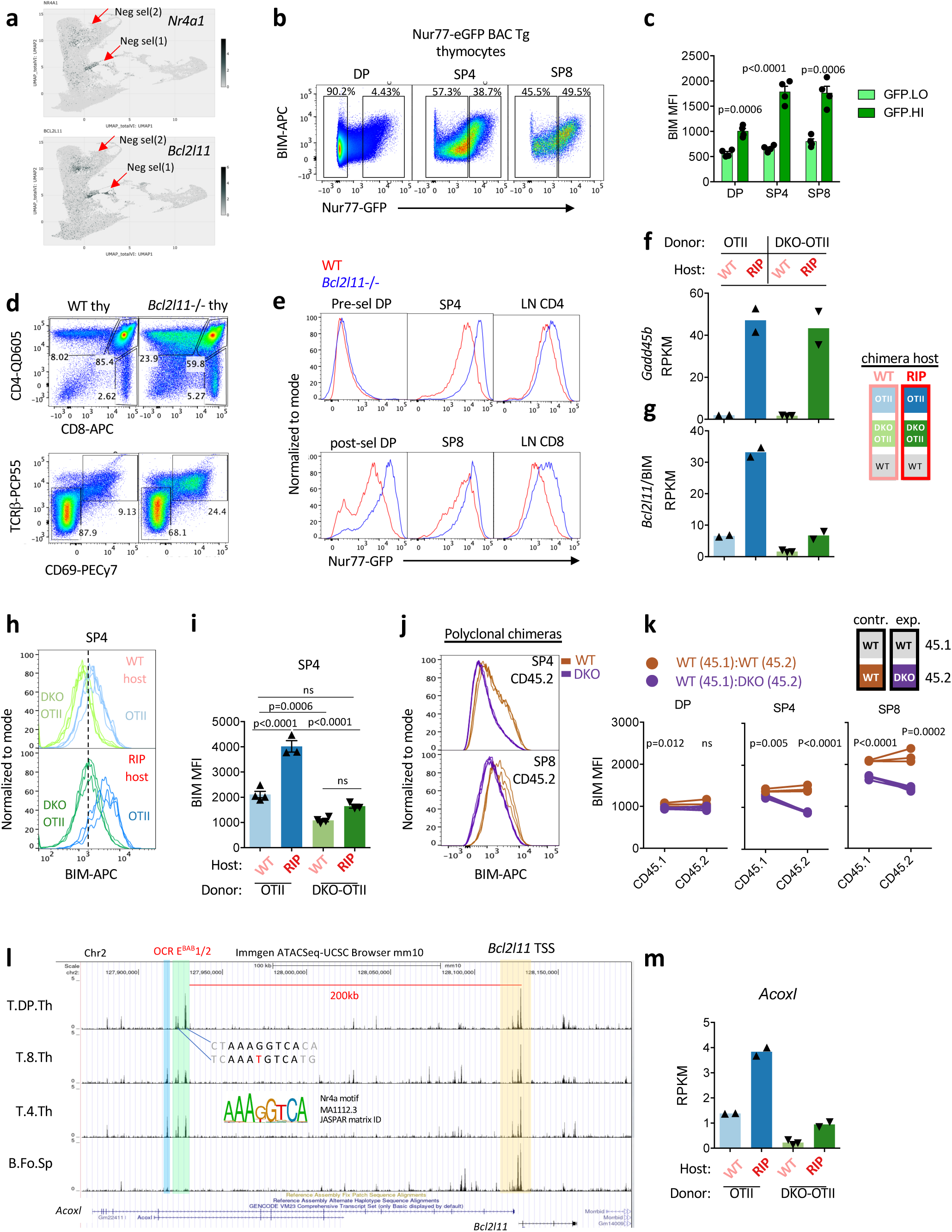
*Nr4a1* and *Nr4a3* are required for thymic transcriptional upregulation of *Bcl2l11*/BIM by RIPmOVA. **a.** *Nr4a1* and *Bcl2l11* transcript UMAP data from publicly available VISION thymus CITEseq interface (http://s133.cs.berkeley.edu:9002/Results.html; Steier et al. Nat Imm. 2023 Sep;24(9):1579-1590.). Red arrows indicate two clusters of negatively selecting cells, early in cortical DP and late in medullary SP. See also Supp Fig 1e for additional annotation. **b, c.** Nur77-eGFP BAC Tg thymocytes were stained to detect thymic subsets and intra-cellular BIM protein. Representative plots depict correlation between markers and show GFPlo/hi gates. Graph (c) depicts MFI of BIM among GFPhi and GFPlo populations as gated in (b) +/– SEM from 4 biological replicates. **d, e.** Nur77-eGFP BAC Tg was crossed onto the *Bcl2l11*–/– background. Thymocytes *Bcl2l11*+/+ and –/– reporter mice were stained to detect SP subsets (d, top panels) and gating for pre-selection (CD69lo TCRβlo) and post-selection (CD69hiTCRβhi) DP thymocytes (d, bottom panels). Histograms depict reporter GFP expression in overlaid genotypes gated as in (d) along with peripheral CD4, CD8 LN T cells. Data are representative of 3 biological replicates. **f, g.** Graphs depict RPKM for *Gadd45b* (f) and *Bcl2l11* (g) in CD69hi Va2+ SP4 thymocytes of either WT-OTII or DKO-OTII donor genotypes from chimeras depicted in 3a. RNAseq was performed on biological replicates (Table 1a depicts p value and fdr for all pairwise comparisons across sample types, GSE235101). **h, i.** Overlaid histograms depict intracellular BIM protein expression in SP4 OTII and DKO-OTII thymocytes from N=4 WT host and N=3 RIPmOVA host chimeras (as schematized in 3a). Graph (i) depicts BIM MFI +/- SEM from these samples. **j, k.** Overlaid histograms depict intracellular BIM protein expression in CD45.2 donor genotypes in SP4 (top) and SP8 (bottom) thymocytes from N=3 control WT:WT chimeras and N=4 experimental DKO:WT polyclonal chimeras (as schematized in 1c). Graph (k) depict BIM MFI across thymic subsets in both CD45.2 and CD45.1 donor genotypes from both types of chimeras. Lines connect samples from the same chimera. **i.** UCSC browser tracks depict ATACseq peaks from public immgen.org data across thymic subsets and Follicular splenic B cells for reference at the *Acoxl/Bcl2l11* locus. 200kb upstream of the TSS of Bcl2l11 are green-tinted ATAC peaks with consensus Nr4a sites that correspond to EBAB enhancer identified in Hojo et al. (PMID 31197149). Blue tint identifies an adjacent peak with striking cell type specificity but only an imperfect Nr4a site. See also Table 1b. **m.** Graph depicts RPKM for *Acoxl* as in f, g, statistical analyses in Table1a. <u>Statistical tests:</u> b, unpaired two-tailed t-test, do not assume equal SD, corrected for multiple comparisons with Holm-Sidak method. i, one way ANOVA with Tukey’s multiple hypothesis test. k, unpaired two tailed t-test, do not assume same SD

To test whether Nr4a family regulates *Bcl2l11* transcription, we sorted semimature CD69hi SP4 Vα2+Vβ5+ OT-II thymocytes of each donor genotype from either WT or RIPmOVA hosts (as in schematic **Fig 3a**) and compared their transcriptomes. RNAseq analysis of OT-II and DKO OT-II semimature thymocytes shows comparable and robust induction of *Gadd45b* (**Fig 4f**, **Table 1a**), a previously identified transcriptional marker of negative selection that is also correlated with *Bcl2l11* transcript in clusters of deleting thymocytes in CITEseq atlas (**Supp Fig 5b**)^59,60^. This indicates that both OT-II and DKO OT-II sense self-antigen in RIPmOVA hosts, and we successfully captured OT-II thymocytes prior to their deletion. By contrast, *Bcl2l11* was induced in OT-II but not in DKO OT-II (**Fig 4g**). Similarly, BIM protein expression is induced in WT OT-II thymocytes from RIPmOVA hosts relative to WT hosts, but not in DKO OT-II cells (**Fig 4h, i**). As in the RIPmOVA model, polyclonal DKO thymocytes also exhibited a large reduction in BIM expression (**Fig 4j, k**). These data argue that DKO thymocytes encounter and respond to self-antigen but fail to upregulate *Bcl2l11* transcript and BIM protein which may account for their escape from deletion.

Because BIM and other BH3-only family members function through stoichiometric sequestration of anti-apoptotic partners, we next assessed transcript abundance of Bcl2 family “executioners” Bax and Bak, their activators (BH3-only pro-apoptotic family members), and their anti-apoptotic inhibitors (**Supp Fig 5c**, **Table 1a**). Among pro-apoptotic members, *Bax* is constitutively expressed and *Bcl2l11*/BIM is induced by RIPmOVA in an Nr4a-dependent manner, but all others are minimally expressed including *Bbc3*/PUMA (despite its genetic implication in this pathway) and *Pmaip1*/NOXA^61^. Of the anti-apoptotic Bcl-2 family members, *Mcl* and *Bcl2* are the most abundant in our dataset but were not induced by antigen stimulation. The pro-apoptotic members *Bcl2a1* and *Bcl2b1* were induced by antigen and correlated to negatively selecting clusters in CITEseq but were Nr4a-independent (**Supp Fig 5b, c**). Of note, *Bcl2* levels are slightly lower in DKO OT-II than OT-II from RIPmOVA hosts (log2FC= -1.172) but much less reduced than *Bcl2l11* (log2FC =-2.533) (**Table 1a, Supp Fig 5c**). This suggests that the stoichiometric balance between pro- and anti-apoptotic family members in DKO thymocytes is shifted to favor survival.

### Conserved Nr4a motifs in an enhancer of thymic *Bcl2l11* transcription

We next sought to determine how Nr4a factors induce *Bcl2l11* transcription. We searched publicly available ATACseq data (Immgen.org) to identify well-defined Nr4a consensus DNA binding motifs in putative cis-regulatory elements near *Bcl2l11*. Peaks at regions of open chromatin (OCR) corresponding to the TSS (transcriptional start site) of *Bcl2l11* did not contain Nr4a consensus motifs. However, we identified a limited number of OCRs within several hundred kb of *Bcl2l11* TSS with highly conserved Nr4a motifs (**Table 1b**). Among these, a pair of peaks within approximately 5kb of one another and 200kb upstream of Bcl2l11 TSS (located within intron 9 of the adjacent gene *Acoxl*) showed uniquely high accessibility in thymocytes and T cell lineage populations but were undetectable in B cells (**Fig 4l, Supp Fig 5d**). Moreover, these two peaks corresponded to a putative enhancer (termed *E^BAB^*) encompassing two co-localized thymus-specific peaks identified on the basis of independent genomic data sets (H3K27ac CHIPseq and DHS) ^62^. CRISPR-Cas9 deletion of this *E^BAB^* cis-regulatory element previously revealed that it is required for thymic induction of *Bcl2l11* transcript and negative selection ^62^. We have therefore termed these two OCRs *E^BAB1^*and *E^BAB2^* and propose that TCR-induced Nr4as bind directly to these enhancers to induce *Bcl2l11* transcription and drive negative selection. Indeed, we identified an *Nr4a1*/Nur77 ChIPseq peak at *E^BAB1^*(GSE102393, Supp Fig 5d)^63^. Strikingly, we found that the adjacent *Acoxl* gene (within which *E^BAB^*sits) similarly exhibited impaired induction in DKO thymocytes suggesting it may be co-regulated with *Bcl2l11* by Nr4as via this cis regulatory element (**Fig 4m**).

**Figure 5.**
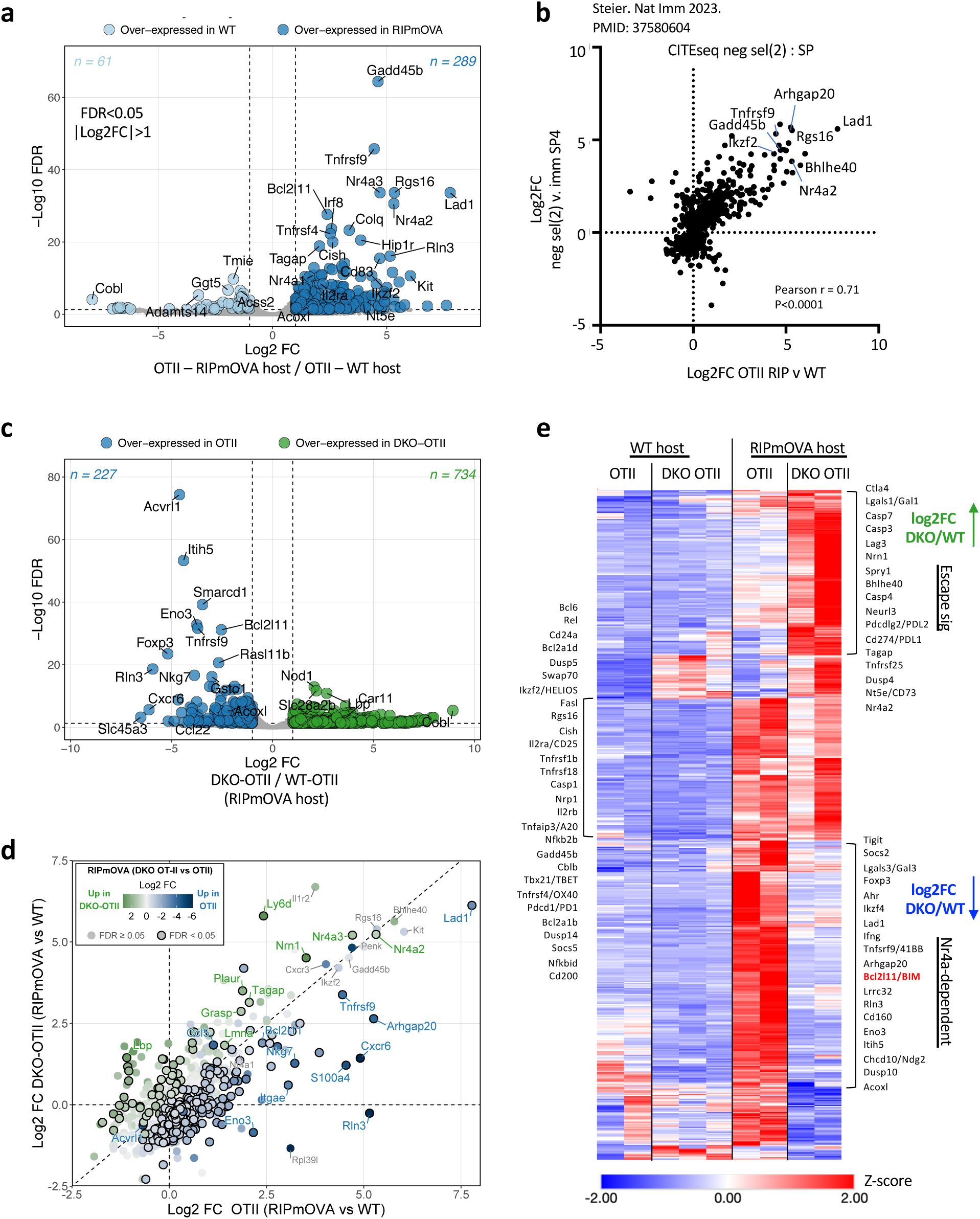
*Nr4a1* and *Nr4a3* mediate a broad transcriptional program induced by antigen recognition in thymic medulla. **a.** RNA sequencing was performed on CD69hi Vα2+ SP4 thymocytes of either WT-OTII or DKO-OTII donor genotypes from host chimeras depicted in 3a. Differential expression analysis was undertaken to identify statistically significant DEG in all pairwise comparisons (Log2FC >1 and fdr <0.05). See also Table 1a, GSE235101. Volcano plot depicts DEG (colored points) in WT-OTII thymocytes sorted from either WT hosts or RIPmOVA hosts. **b.** FC FC plot compares OTII chimera RNAseq data set described in (a) with VISION CITEseq data. Graph depict log2FC OTII thymocytes from RIPmOVA/WT hosts and log2FC neg sel(2) cluster / immature SP4 cluster from VISION thymus CITEseq. Pearson correlation coefficient is quantified. See also Table 2c. **c.** As in (a) except volcano plot depicts DEG (colored points) in WT-OTII vs. DKO-OTII thymocytes sorted from RIPmOVA hosts. **d.** Log2FC for OTII thymocytes from RIP/WT hosts plotted against log2FC for DKO-OTII thymocytes from RIP/WT hosts. Genes with RPKM > 5 for all samples in at least one experimental group are plotted. Superimposed color scheme identifies DEG for comparison of DKO-OTII and WT-OTII SP thymocytes from RIPmOVA hosts (as in c.) Superimposed colored points represent Log2FC comparing DKO-OTII and WT-OTII from RIPmOVA hosts and outlined points indicate fdr < 0.05 for this comparison. **e.** Heatmap depicts all pairwise DEG (Log2FC >1 and fdr <0.05) between any of the 4 populations from data set described in (a) that are also upregulated (log2FC > 0) in WT-OTII thymocytes from RIPmOVA hosts relative to WT hosts (N=551; Table 1a). Annotated modules depict selected genes that are upregulated, downregulated, or comparably expressed in DKO-OTII > WT-OTII from RIPmOVA hosts.

### Self-antigen encounter in the medulla induces a transcriptional program associated with negative selection

We next took advantage of our model to define the global transcriptional response of SP thymocytes encountering model self-antigen expressed in mTECs (chimera schematic **Fig 3a**). 350 transcripts are differentially expressed in WT OT-II semimature SP4 thymocytes sorted from WT and RIPmOVA hosts (|log2FC|>1, FDR < 0.05; **Fig 5a**; **Table 1a**). Of these, the majority are upregulated, and only a small number of genes are downregulated. This program is highly conserved (82/350 transcripts) across the related model system in which model antigen hen egg lysozyme (HEL) is expressed under the control of the insulin promoter (InsHEL) together with the 3A9 HEL-specific TCR Tg, but minimally shared with the HY-CD4 model in which ubiquitous self-antigen deletes cognate TCR Tg cells early at the DP stage in the cortex (**Supp Fig 6a, b**, **Tables 2a, b**)^34,60^. Notably, *Nr4a1, Nr4a3*, and *Bcl2l11* are among the limited set of shared genes induced in each of these models.

We next compared antigen-induced transcripts in our bulk RNAseq data with the set of transcripts enriched in *Bcl2l11*+ CITEseq clusters fated for negative selection (**Fig 4a**). As noted earlier, two deleting clusters are identified, one corresponding to CCR7-CD69+ DP thymocytes likely in the cortex, and the second corresponding to CCR7+ CD69hi semimature SP thymocytes likely encountering Ag in the medulla (**Supp Fig 1e**). Correlation between our OT-II/RIPmOVA data and the medullary deleting cluster is remarkably high, again suggesting a highly conserved transcriptional program is triggered by self-antigen recognition in the medulla (**Fig 5b**, **Table 2c**). By contrast, the transcriptional programs in cortical DP thymocytes and in medullary SP thymocytes undergoing negative selection diverge; only a limited number of DEG are shared between TCR Tg models and both deleting clusters, including the *Nr4a* genes and *Bcl2l11*/BIM (**Supp Figs 6c, e, Table 2d**).

Importantly, the cluster of medullary thymocytes undergoing negative selection are adjacent to tTreg on the CITEseq UMAP, highlighting their transcriptional similarity to one another (**Supp Fig 6e-g**, **Table 2d**). While *Bcl2l11* and *Gadd45b* are higher in cells fated for deletion than in Treg, several transcripts upregulated in response to RIPmOVA set mark both of these populations, such as the transcription factor *Ikzf2*/HELIOS, the co-stimulatory receptors *Tnfrsf4*/OX40, *Tnfsrf9*/41BB, as well as *Nr4a1* and *Nr4a3* themselves (**Supp Fig 6e, f**). This is consistent with establishment of a common precursor pool of self-reactive thymocytes ^45^. Importantly, our sorted OT-II samples express virtually no detectable *Foxp3* transcript and also have rather low *Il2ra* levels suggesting that profiled thymocytes were captured before commitment to either deletion or diversion and are distinct from CD25hi pre-Treg^45^, consistent with flow cytometric analyses (**Supp Fig 6g**, **Fig 3d-f**).

### Nr4a1 and Nr4a3 are required to enforce a broad SP thymocyte transcriptional program in response to self-antigen recognition in the medulla

We next sought to determine how the large and conserved transcriptional program induced by antigen recognition in the thymus was regulated by Nr4as. Principal component analysis revealed that DKO OT-II and WT OT-II thymocytes diverged markedly in RIPmOVA (but not WT) hosts, suggesting Nr4as play a selective role in self-reactive thymocytes (**Supp Fig 7a, b**, **Fig 5c**). We identified genes that were differentially expressed in Nr4a-sufficient and deficient OT-II thymocytes from RIPmOVA hosts (**Fig 5c, d**). To visualize this program across both donor and host, we identified transcripts differentially expressed between any of the 4 sorted thymocytes populations (fdr < 0.05, |log2FC| >1), and generated a heatmap focusing on those that are upregulated by antigen recognition (log2FC>0 OT-II-RIP/OT-II-WT, N=551) (**Fig 5e**, **Table 1a**). A set of RIPmOVA-induced transcripts is shared between WT OT-II and DKO OT-II thymocytes (**Fig 5a, d, e, Supp Fig 7c**). These include *Gadd45b* (as noted earlier), *Ikzf2*/HELIOS as well as inhibitory receptors *Pdcd1*/PD1, *Cd200*, co-stimulatory receptor *Tnfrsf4*/OX40, and the ubiquitin ligases *Cblb* and *Tnfaip3*/A20. These data confirm that there is no global defect in antigen sensing or TCR signaling in DKO thymocytes.

Notably, over one third of the RIPmOVA-induced transcriptome is almost entirely lost in the absence of Nr4as (**Fig 5e**). This was similar to a broad - but not global - transcriptional defect previously described in the autoimmune-prone NOD genetic background that was also associated with impaired *Bcl2l11* induction and defective negative selection (**Supp Fig 7d**, **Table 2a**)^35,60^. The Nr4a-dependent program we identify encompasses Treg-associated transcripts (e.g. *Foxp3* itself despite minimal reads, *Ikzf4, Ahr, Lrrc32*/GARP), and negative selection-associated transcripts *Lad1, Arhgap20*, as well as *Bcl2l11*/BIM itself and the co-regulated gene *Acoxl*, as shown earlier (**Figs 5c, d, e**). We also identified Nr4a-dependent transcripts *Rln3, Eno3, Itih5* and *Tnfrsf9*/41BB previously reported in thymocytes and peripheral T cells, including *Acvrl* which is adjacent to *Nr4a1* in the mouse genome (**Figs 5c-e, Supp Fig 7b, e, f, Supp Fig 8a, b, Tables 2e-i)^16,64^**. We also confirmed Nr4a-dependence of Ndg2/Chchd10 – a Nur77-induced transcript with pro-apoptotic functions identified by Winoto and colleagues, but we find that

other putative Nur77 targets reported in that study are Nr4a-independent (*Fasl, Pdcd1/*PD1, TRAIL/*Tnfsf10*, *Ctla4*, and Ndg1/*Khdc1*) ^65^.

### Self-antigen encounter in the medulla induces an anergy-like transcriptional program that is partially dependent on Nr4as

We next sought to understand the functional significance of this Ag-dependent thymic transcriptional program by comparing it to the transcriptome of mature T cells under different conditions. Many genes upregulated by RIPmOVA in SP thymocytes were not specifically associated with survival/apoptosis (**Fig 5a, e**, **Table 1a**). When we compare these genes to TCR-dependent transcripts induced in peripheral T cells, we noted that only a minority represent rapidly induced primary/early response genes, while the bulk of the self-antigen-induced transcriptome in medullary SP thymocytes strongly resembles that of naturally occurring anergic T cells (**Fig 6a**, **Table 2i**).

**Figure 6.**
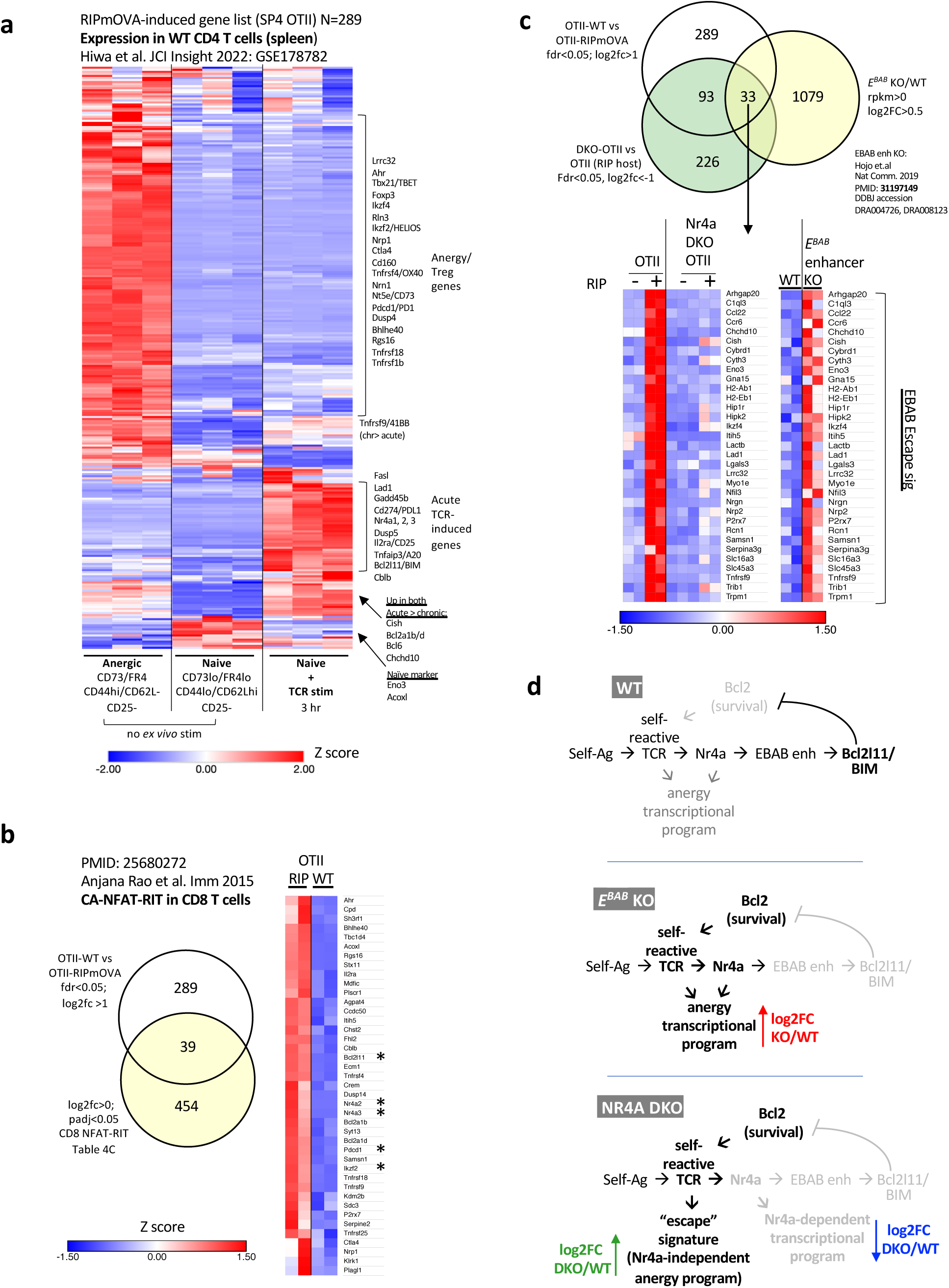
Self-antigen encounter in the medulla induces an anergy-like transcriptional program that is partially dependent on Nr4as. **a.** Heatmap depicts dataset GSE178782 encompassing transcriptome of naïve (CD62Lhi CD44lo FR4lo CD73lo CD25-) CD4 T cells of WT polyclonal genotype +/- 3hr anti-CD3 stimulation as well as ex vivo naturally occurring anergic CD4 T cells (CD62Llo CD44hi FR4hi CD73hi CD25-). Genes included in the heatmap are those identified in our data set (Fig 5a) that are DEG in WT-OTII thymocytes in RIPmOVA hosts relative to WT hosts (log2FC > 1, fdr< 0.05, N=286). Modules depict selected genes that are either primary response genes, anergy-associated genes, or naïve markers. See also Table 2i. **b.** Venn diagram depicts overlapping genes between thymic DEG identified in our data set (OTII from RIPmOVA v WT hosts) and transcripts induced in CD8 T cells by expression of CD8NFAT-RIT (PMID: 25680272). Heatmap depicts overlapped gene expression in OTII thymocytes from RIPmOVA or WT hosts. See also Table 2j. **c.** Venn diagram depicts RIPmOVA-induced genes that are Nr4a dependent (white/green circle overlap) among genes overinduced in EBAB enhancer KO thymocytes (yellow circle). Heatmaps depict expression of those genes in our thymic RNAseq data set aligned against expression in WT or EBAB KO RNAseq data set (PMID: 31197149; see also Table 2m) **d.** models schematize regulation of Ag-dependent transcriptional modules in thymocytes that give an over-induced “escape” signature among either EBAB KO or Nr4a1/3 DKO cells that escape BIM-induced deletion. The escape signature is uncoupled from Nr4a-dependent program in DKO cells.

Anergic gene expression is induced when T cells encounter signal 1 (TCR) in the absence of signal 2 (CD28 co-stimulation). Consistent with this, RIPmOVA-induced thymic transcripts overlap with a CD4 T cell tolerance program induced in vitro by TCR stimulation without co-stimulation (**Supp Fig 8c**, **Table 2j**) ^66^. Signal 1 in the absence of Signal 2 produces NFAT nuclear translocation in the absence of AP-1 and NF-kB^67^. Anjana Rao and colleagues showed that a constitutively active form of NFAT that cannot cooperate with AP-1 (CA-NFAT-RIT) drives an anergic transcriptional program^68^. Here again we identified a set of NFAT-responsive genes that overlapped with our data set (**Fig 6b**, **Supp Fig 8d**, **Table 2k**) and account for some of the anergy and exhaustion programs discovered in peripheral T cells such as *Nr4a2/3, Tnfrsf4* and *9*, as well as *Cblb* and *Bhlhe40* among others. Similarly, markers of bona fide CD8 T cell exhaustion are shared with OT-II thymocytes from RIPmOVA hosts (**Supp Fig 8e**, **Table 2l**)^69^.

Importantly, many anergy-associated genes induced by RIPmOVA required Nr4a for their induction (**Fig 5e**). We discovered negative regulators of TCR and cytokine signaling in this module, including cell surface molecules *Tigit, Cd160, Lgals3*/Galectin-3, as well as intracellular suppressor of cytokine signaling *Socs2* and MAPK PTPase *Dusp10*. Reducing stringency of filtering (p<0.01) to increase discovery of RIPmOVA-induced genes also identifies *Cd5, Dusp5, Dusp1, Tox2*, and *Cish* as negative regulators modulated by the Nr4a family (**Table 1a**, GSE235101).

### Nr4a-deficient thymocytes exhibit a transcriptional escape signature

In addition to identifying genes that depend on Nr4as for their induction, we observed a module of antigen-induced transcripts in OT-II thymocytes that were over-expressed in the absence of Nr4as (**Fig 5e**). We hypothesize that these genes largely represent an “escape signature” in DKO SP4 thymocytes that were destined for - but evaded - deletion; over-induced genes include those that are upregulated by strong TCR signaling but do not require Nr4as for their induction.

Indeed, a similar “escape signature” was described among thymocytes evading negative selection in *E^BAB^* KO mice harboring deletion of a thymic *Bcl2l11*/BIM enhancer (**Fig 6c, d**, **Table 2m**)^62^. Strikingly, some (N=33) of this TCR-dependent escape signature was Nr4a-dependent including *Arhgap20, Lad1, Tnfrsf9, Eno3* among others. This is consistent with our model that Nr4as are upstream of the *E^BAB^* enhancer to induce *Bcl2l11*.

By contrast, the escape signature in DKO thymocytes represent TCR-dependent genes that do not require Nr4as for their induction (**Fig 6d**, schematic model), and includes many anergy-associated inhibitory cell surface molecules: *Ctla4, Lgals1*/Galectin-1, genes encoding both PDL1 (*Cd274*) and PDL2 (*Pdcdlg2*), and *Lag3* as well as the anergy-associated marker *Nt5e*/CD73 which is expressed on naturally-occurring anergic populations (**Fig 5e**)^70^. Additional negative regulators and anergy-associated genes in this module included *Dusp4*, the E3 ubiquitin ligase *Neurl3*, the MAPK negative regulator *Spry1*, the NFAT-induced TF *Bhlhe40*, as well as caspases *Casp3/4/7* (**Fig 5e**).

### Epigenetic imprint of thymic antigen encounter persists in peripheral RTEs with “anergic” phenotype

We (Au-yeung and colleagues) recently described a population of naturally occurring naïve CD4 T cells (CD62Lhi CD44lo CD25-Foxp3-) characterized by high Nur77-eGFP and low Ly6C expression (**Fig 7a**). By contrast to other naïve CD4 T cells, these Ly6CloGFPhi cells (termed population “D”) express inhibitory molecules, exhibit impaired response to TCR stimulation, and preferentially differentiate into Treg ^71,72^. Ly6C is developmentally regulated with very low expression in the thymus that increases in the periphery early after thymic egress (**Fig 7a, b**)^73^. Indeed, we found that population D was highly enriched for recent thymic emigrants (RTE) based on markers Qa2 and CD24 (**Fig 7b, c, d**) ^74^.

**Figure 7.**
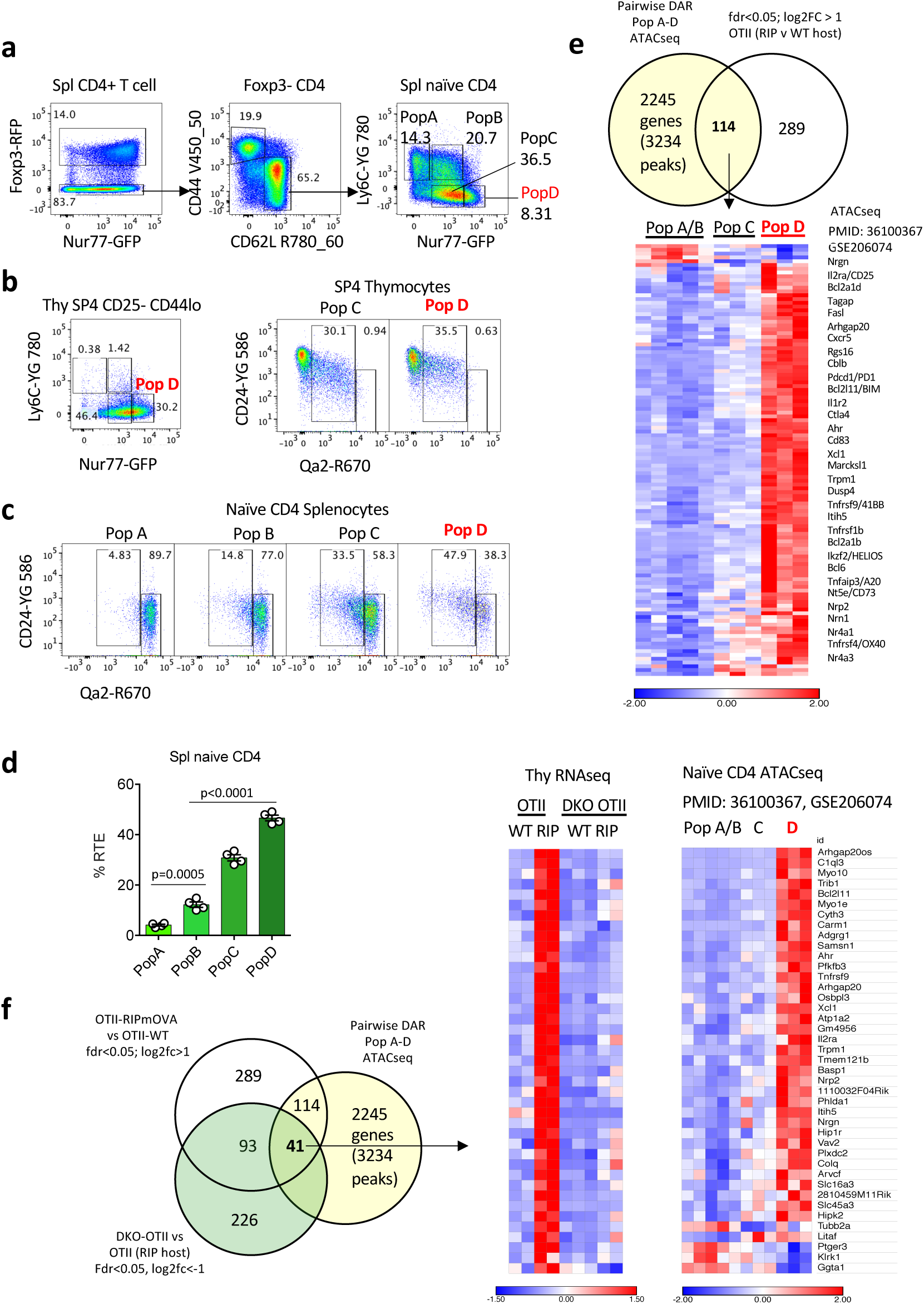
Epigenetic imprint of thymic antigen encounter persists in peripheral RTEs with “anergic” phenotype. **a.** Gating scheme to identify Pop A, B, C, D among naïve CD4 splenic T cells from mice harboring both Foxp3-RFP reporter and Nur77-eGFP reporter. Sequential gating excludes Foxp3+ Treg and defines Pop A-D among naïve CD62Lhi CD44lo T cells. **b.** Representative plot (left) depict analogous Pop A-D gates as shown in panel (a) among CD25-SP4 thymocytes. Representative plots (right) depict maturity markers CD24 and Qa2 among pop C, D SP4 thymocytes as gated on left. **c, d.** Representative plots depict maturity/RTE markers CD24/Qa2 (stained in parallel with samples in b) among splenic naïve CD4 Pop A-D as gated in panel (a). Graph (d) depicts quantification of % RTE (CD24hi Qa2lo) among Pops A-D as gated in (c) from 4 biological replicates +/- SEM. **e.** Venn diagram depicts overlap between RIPmOVA-induced transcriptional program and genes near differentially accessible regions (DAR) in the genome across Pops A-D (PMID: 36100367; GSE206074). To do so, DAR were collapsed by gene and aligned with OTII/RIPmOVA RNAseq data set. Heatmap depicts rpm in Pops A-D for most differentially accessible OCR near these genes (see also Table 2n). **f.** Venn diagram as in (e) but further filtered to identify Nr4a-dependent gene set. Heatmap depicts expression of those genes in our thymic RNAseq data set aligned against accessibility in Pop A-D ATACseq (PMID: 36100367; GSE206074; see also Table 2n) <u>Statistical analysis:</u> d, One way Anova with Tukey’s data in a-d are representative of at least N=4 biological replicates.

ATACseq analysis revealed a marked increase in chromatin accessibility across a set of approximately 2000 genes in population D relative to other naïve CD4 T cells and this signature correlated with RTE enrichment among naïve T cell populations, suggesting a distinct history potentially reflecting recent thymic signaling (**Fig 7d, e**) ^71^. These include genes associated with anergy and exhaustion (**Fig 7e**). We compared these DARs to the set of genes upregulated in OT-II thymocytes from RIPmOVA hosts and observed a striking enrichment; almost half of all RIP-induced genes exhibited greater chromatin accessibility in popD/RTEs (**Fig 7e**, **Table 2n**). Moreover, among the overlapping genes, nearly half require Nr4as for their induction (49/114; **Fig 7f**). Together, these findings led us to postulate recent antigen-encounter in the thymus left an Nr4a-dependent epigenetic imprint on Pop D that confers an “anergy”-like state in the periphery.

### Nr4a-deficient thymocytes acquire an antigen-dependent anergic program in the thymus

We next sought to identify DKO thymocytes fated for - but rescued from - negative selection. We focused on RIPmOVA-induced transcripts that do not require Nr4as for their induction and therefore could mark strongly signaled thymocytes escaping deletion (**Fig 8a-d**). HELIOS is encoded by *Ikzf2* and - in addition to high expression among Treg - was previously identified as a marker of Foxp3-negative thymocytes escaping negative selection ^37^. Indeed, as noted earlier, *Ikzf2* transcript is upregulated by RIPmOVA in our bulk data set, and it is also highly expressed in both negatively selecting clusters in CITEseq data (**Fig 8a, b**). We find that *Ikzf2* is upregulated in both OT-II and DKO OT-II SP4 in RIPmOVA hosts. HELIOS protein expression is high (comparable to Treg) in a large subset of these cells in the negatively selecting hosts but virtually undetectable in WT hosts (**Fig 8e, f**). Flow cytometric analysis of protein expression reveals a larger fraction and much larger total number of DKO OT-II SP4 thymocytes are HELIOS positive (**Fig 8e-g**). We propose these Foxp3-cell represent DKO thymocytes that have escaped negative selection.

**Figure 8.**
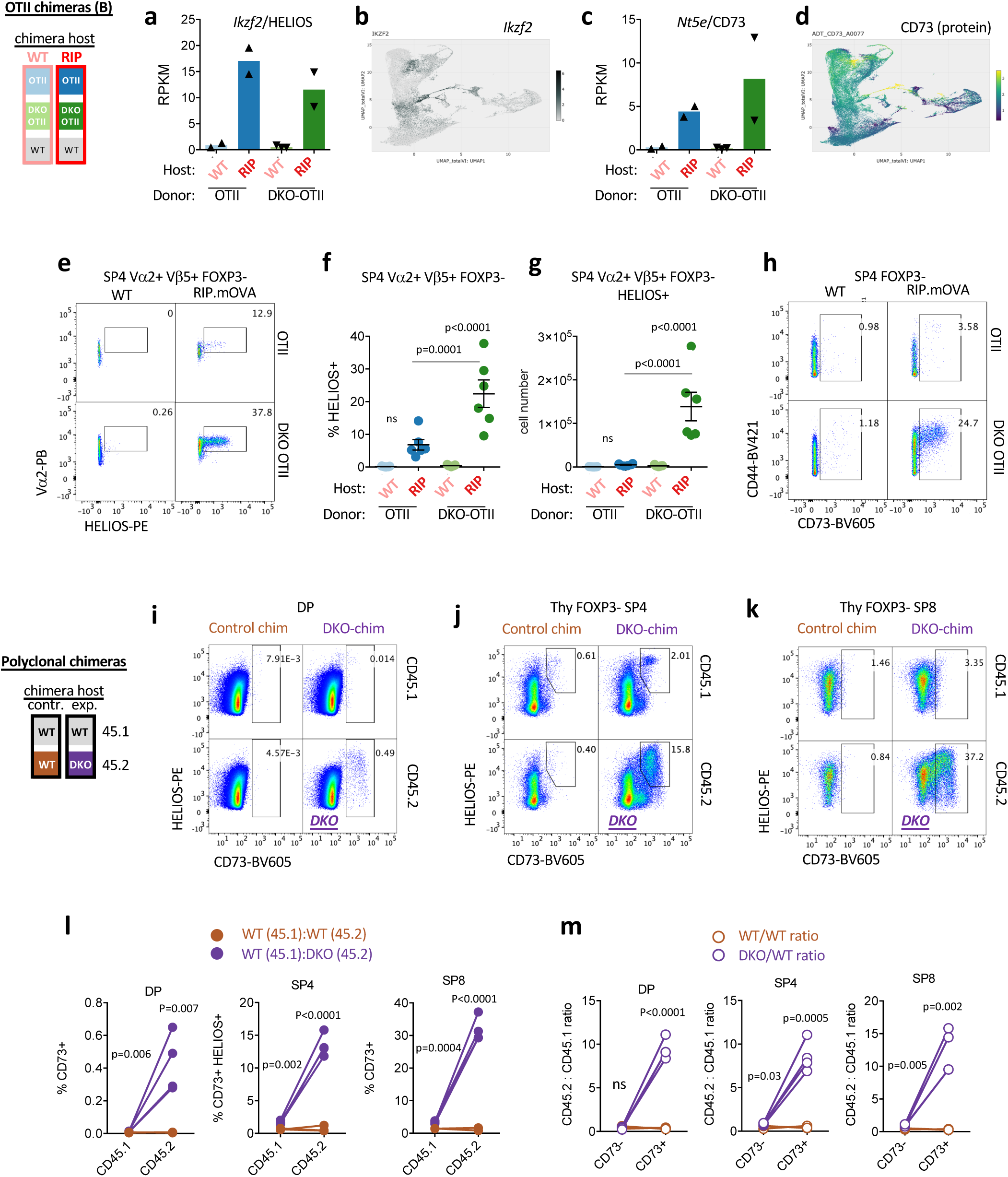
Nr4a-deficient thymocytes acquire an Ag-dependent anergic program in the thymus. **a-d.** Panels depict Ikzf (a, b) and Nt5e/CD73 (c, d) expression our RNAseq data (a, c) and VISION CITEseq (b, d) data sets as described earlier. RPKM graphs show transcript abundance in CD69hi Va2+ SP4 thymocytes of either WT-OTII or DKO-OTII donor genotypes from chimeras depicted in 3a. (Table 1a depicts p value and fdr for all pairwise comparisons across sample types, GSE235101). UMAP data (b, d) from publicly available VISION thymus CITEseq interface (http://s133.cs.berkeley.edu:9002/Results.html; Steier et al. Nat Imm. 2023 Sep;24(9):1579-1590.) **e-g.** Representative plots (e) depict gating to identify HELIOS+ cells among SP4 Foxp3-OTII thymocytes of each genotype from RIPmOVA and WT host chimeras (see 3a schematic). Graphs depict % (f) and absolute number (g) of HELIOS+ cells as gated in (e). **h.** Representative plots depict gating to identify CD73+ cells among SP4 Foxp3-OTII thymocytes of each genotype from RIPmOVA and WT host chimeras (see 3a schematic). Quantification is shown in Supp Fig 9a, b. **i-k.** Representative plots depict gating among DP (i), SP4 (j), and SP8 (k) subsets to identify HELIOS+CD73+ cells of each donor genotype from experimental DKO:WT polyclonal chimeras and from control WT:WT polyclonal chimeras (see 1c schematic). **l.** Quantification of % CD73/HELIOS gates shown in i-k from each donor genotype in panels i-k. Lines connect donors in the same chimera. **m.** Quantification of ratio of donor genotypes CD45.2/CD45.1 from each chimera among CD73/HELIOS in panels i-k. <u>Statistical tests:</u> f, one way ANOVA with pre-specified comparisons corrected for by Sidak test g, one way ANOVA with Tukey’s multiple hypothesis test. l, m, unpaired two-tailed t-test, do not assume equal SD Graphs in f, g, depict N=7 WT host and N=6 RIPmOVA host chimeras. Graphs in l, m depict N=3 WT:WT control chimeras and N=4 DKO:WT chimeras. representative of at least 3 independent sets of chimeras.

Among transcripts enriched in DKO OT-II cells relative to WT OT-II from RIPmOVA hosts (so-called “escape signature”) is *Nt5e* which encodes CD73 (**Fig 8c**). CD73 is a marker of naturally occurring CD4 anergic cells together with FR4 ^70^.As with HELIOS, CD73 is induced in a large subpopulation (>20%) of DKO OT-II SP4 (Foxp3-neg) only in RIPmOVA but not WT hosts (**Fig 8h, Supp Fig 9a, b**). By contrast, the analogous WT OT-II cell population in RIPmOVA hosts is largely absent, consistent with their deletion in RIPmOVA.

Similar to RIPmOVA model, polyclonal Foxp3-neg DKO SP4 also harbor a large (>20%) CD73hi population that are also HELIOS+ (**Fig 8i-m**). This compartment far exceeds the size of a WT tTeg population and is apparent despite a diverse repertoire of T cell clones and endogenous antigens. We postulate that the appearance of this population reflects escape of DKO cells from negative selection by endogenous antigen recognition. Strikingly, among polyclonal DKO SP8 thymocytes, CD73hi population was also uniquely expanded and constituted nearly half of all SP8s (**Fig 8k, l, m**). This suggested that a larger fraction of SP8 than SP4 are rescued from deletion, consistent with a more stringent threshold for deletion in that lineage^43,44^. This moreover suggests that altered SP4/SP8 ratio in DKO reflects differential threshold for deletion rather than altered lineage specification. Importantly, both OT-II and polyclonal DKO CD73hi cells express CD24 and CD62L demonstrating that they are indeed bona fide thymocytes rather than recirculating mature T cells^40,74^.

### DKO thymocytes exhibit an anergic imprint that persists in the periphery

Consistent with thymic deletion, WT OT-II CD4 splenic T cells are almost entirely lost in RIPmOVA hosts (**Fig 9a, Supp Fig 9c**). Strikingly, not only do DKO OT-II CD4 splenic T cells persist, but they include a large CD73hi sub-population that is evident early after reconstitution (**Fig 9a-d**) and corresponds to a similar population concurrently detected among DKO OT-II SP4 thymocytes (**Fig 8e-h, Supp Fig 9a, b**), suggesting they may represent the same cells captured after thymic egress. Because this phenotype is evident in splenic naïve T cells while RIPmOVA is tissue-restricted, it is possible this reflects thymic rather than peripheral self-antigen encounter.

**Figure 9.**
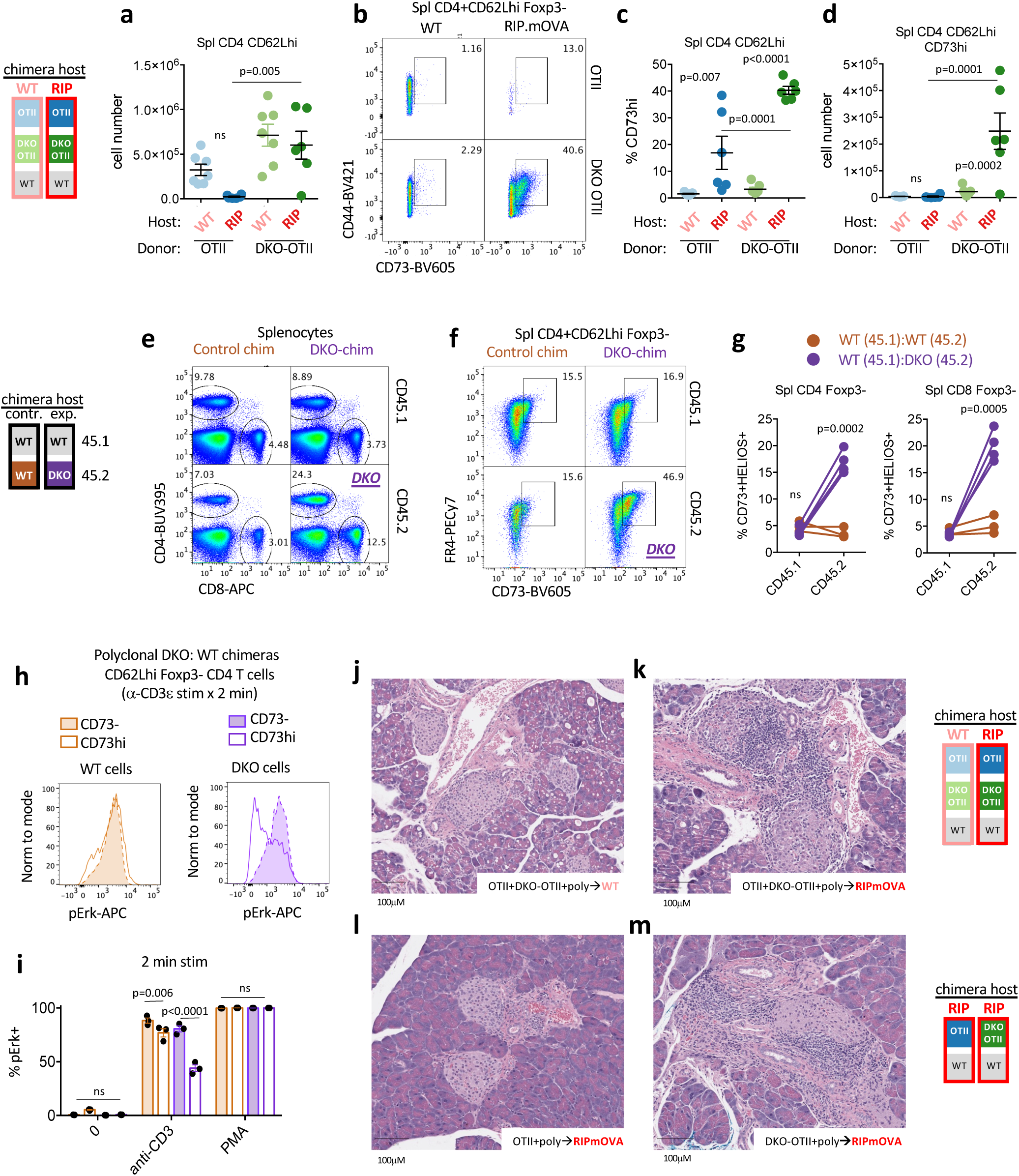
DKO T cells acquire an anergic imprint in the thymus that persists in the periphery, but break tolerance. **a.** Graph depicts naïve CD4 splenic T cell count of each donor genotype (OTII or DKO-OTII) from WT or RIPmOVA host chimeras (see Fig 3a schematic). **b-d.** Representative plots depict gating to identify CD73+ cells among Foxp3-CD4 naïve T cells of each donor genotype (OTII or DKO-OTII) from RIPmOVA and WT host chimeras (see 3a schematic). Graphs depict % (c) and absolute number (d) of CD73+ cells as gated in (b). **e.** Representative plots depict gating to identify splenic CD4 and CD8 T cells of each donor genotype from polyclonal chimeras (see Fig 1c schematic). See supp Fig 9g for quantification. **f.** Representative plots depict gating to identify CD73+ cells among Foxp3-CD4 naïve T cells of each donor genotype from polyclonal chimeras (see Fig 1c schematic). **g.** Graphs depict % CD73+/HELIOS+ cells as gated in Supp fig 9f. from each donor genotype in polyclonal chimeras (see Fig 1c schematic). Lines depict donor genotypes from the same chimera. **h, i**. Splenocytes from polyclonal DKO:WT chimeras (as in schematic Fig 1c except 14 week reconstitution) were stimulated with anti-CD3/cross-linking Ab for 2 minutes, followed by fixation, permeabilization and staining to identify intracellular pErk as well as congenic markers CD45.2/CD45.1, Foxp3, CD44, CD62Lhi, CD4, and CD73/FR4 to identify naïve Foxp3-CD4 T cells with either high or low CD73 expression of each donor genotype. Representative overlaid histograms depict pErk in these population from either CD45.1 WT (brown) or CD45.2 DKO (purple) cells. Graph depicts quantification of pErk+ gate across these populations from N=3 biological replicates +/- SEM. **j, k.** representative histology from WT (j) or RIPmOVA (k) host chimeras containing donor OTII and DKO-OTII and polyclonal BM as schematized (see also Fig 3a chimera design) at 6 weeks following irradiation/BM transfer. **l, m.** representative H&E pancreas histology from RIPmOVA host chimeras containing either WT-OTII:poly or DKO-OTII:poly donor BM as schematized (see Fig 2a chimera design) at 10.5 weeks following irradiation/BM transfer. <u>Statistical tests:</u> a, c, d, one way ANOVA with Tukey’s multiple hypothesis test. Graphs in a, c, d, depict N=7 WT host and N=6 RIPmOVA host chimeras. g, unpaired two-tailed t-test, do not assume equal SD Graphs in g depict N=3 WT:WT control chimeras and N=4 DKO:WT chimeras. representative of at least 3 independent sets of chimeras. i. ordinary two-way ANOVA with Tukey’s multiple comparisons test. Data in j, k are representative of 7, 6 biological replicates respectively. Data in l, m are representative of 3 biological replicates each.

DKO T cells from polyclonal chimeras also accumulate relative to WT in the periphery of competitive chimeras and express anergy-associated phenotypic markers at an early time point in similar proportion to DKO thymocytes (**Fig 9e-g**, **Fig 8j-m, Supp Fig 9d-i**). These polyclonal T cells are CD73hi, co-express FR4, and a subset are also HELIOS+ (**Fig 9f, Supp Fig 9f, h**).

Importantly, DKO CD73hi HELIOS+ T cells from both polyclonal and OT-II chimeras are Foxp3-negative and exhibit a naïve CD62Lhi CD44lo surface phenotype (; **Supp Fig 9e**), similar to PopD (see **Fig 7)**^71,72^ but distinct from a previously described CD44hi anergic population ^70^. Moreover, these cells not only exhibit CD73hi, FR4hi phenotype, but are also functionally anergic – as evidenced by impaired induction of pErk following in vitro TCR stimulation (**Fig 9h**).We propose that this cell-intrinsic, anergy-like state is an alternate or fail-safe fate for self-reactive DKO thymocytes that evade deletion and diversion, marked by CD73 and HELIOS expression, partially Nr4a-dependent, induced in the thymus, and persistent in the periphery.

### Nr4a required to maintain tolerance to TSA

RIPmOVA - but not WT - host chimeras harboring both OT-II and DKO OT-II donors (schematic **Figure 3a**) develop elevated blood sugar consistent with diabetes (**Supp Fig 9j**).

Indeed, representative pancreatic sections reveal islet immune infiltration and, in some cases, frank destruction (**Fig 9j, k**). Because WT OT-II T cells are efficiently deleted in the thymus while DKO OT-II T cells escape to the periphery and persist in RIPmOVA hosts (**Fig 9a**), we hypothesize that DKO OT-II peripheral T cells promote disease in these chimeras. Importantly, the majority of peripheral Treg in these chimeras – normally reconstituted - are of polyclonal donor origin as seen during thymic development (**Fig 3c-f, Supp Fig 9c**). While the small remaining number of OT-II splenic CD4 T cells in RIPmOVA hosts are largely diverted to Foxp3+ fate in periphery (30% of surviving cells), this is not true for DKO OT-II peripheral T cells, all of which are Foxp3-neg, even in RIPmOVA (**Supp Fig 9c**).

To isolate contribution of WT OT-II and DKO OT-II cells in disease pathogenesis, we sought to track them in separate hosts. To this end, we returned to an earlier set of chimeras in which either WT OT-II or DKO OT-II donors were mixed with excess of polyclonal BM. This mixture was transferred into two distinct set of irradiated RIPmOVA hosts (see schematic, **Figure 2a**). As described earlier, WT OT-II cells in RIPmOVA recipients were counter-selected relative to polyclonal T cells and the few remaining cells were largely diverted into the Treg compartment, while DKO OT-II were neither deleted nor diverted (**Fig 2d-i**). Finally, we found again islet inflammation in RIPmOVA recipients of DKO OT-II but not WT OT-II cells (consistent with lack of disease in prior reports^51^), suggesting Nr4as are required to maintain tolerance to the model TRA RIPmOVA (**Fig 9l, m, Supp Fig 9k**).

## Discussion

Deletion of self-reactive thymocytes is a vital central T cell tolerance mehcanism, yet the molecular pathways that impose this fate have been elusive. Here we identify members of the Nr4a family as the essential factors that link self-antigen recognition to a broad transcriptional program that imposes not only negative selection and Treg diversion, but also a cell-intrinsic anergy-like tolerance program in developing thymocytes.

Although *Nr4a1* and *Nr4a3* exhibit functional redundancy, their regulation and expression diverge in critical ways. Prior studies suggest that the half-life of Nr4a1/Nur77 protein is shorter than that of Nr4a3/Nor1^23,75^(immpres.co.uk). This may help explain functional redundancy with *Nr4a1* despite low *Nr4a3* transcript abundance. Indeed, Nr4a3 SKO mice exhibit a partial defect in negative selection of OTI thymocytes by RIPmOVA ^17^. The rapid turnover of Nr4a1/Nur77 protein^23,26,27^ may prevent substantial accumulation of Nr4a1/Nur77 protein following the brief, intermittent TCR signals during positive selection. However, Nr4a1 accumulation is more likely following sustained TCR-APC contacts (>3 hrs) shown to be required for deletion ^30,76^. Induction of *Nr4a2* and *Nr4a3* is exclusively dependent on NFAT and exhibits a higher TCR signaling threshold compared to *Nr4a1* ^77^. Expression of *Nr4a3* in addition to *Nr4a1* may tip the balance towards deletion in response to strong TCR signaling. We postulate that Nr4as may function as high pass filters to translate only sustained, high affinity pMHC interactions into tolerogenic fates.

Nr4as have long been implicated in Ag-induced death of lymphocytes, but the mechanism remains controversial. In a proposed post-translational mechanism, Nur77/Nr4a1 induces a conformational change in Bcl2 that exposes its BH3-only domain, thereby transforming this anti-apoptotic factor to a “killer”^78^. This shifts the balance between pro- and anti-apoptotic Bcl2 family members, triggering caspase-induced apoptosis. However, evidence for this pathway during thymic negative selection is circumstantial and limited to DP thymocytes ^79,80^. A transcriptional mechanism was alternatively proposed because truncated Nr4a1 constructs that retain their DNA-binding domain rescue deletion and suppress transcription by other Nr4a family members in a dominant negative manner ^10,14,81^. Here we show Nr4as function definitively in a transcriptional (“genomic”) capacity to regulate *Bcl2l11*/BIM transcription in self-reactive SP thymocytes^56^.

*Bcl2l11*/BIM and *Nr4a*/Nur77 are coordinately upregulated first in strongly signaled post-selection DP thymocytes in the cortex by excessive self pMHC reactivity. BIM has been implicated in an early wave of deletion at this stage, although this role varies according to model system ^9,57,61^. Dominant negative Nur77/Nr4a1 Tg constructs were reported to interfere with deletion driven by super Ag and ubiquitous model Ag ^13,81^. We observe only a subtle initial selective advantage for polyclonal DKO thymocytes in the post-selection DP compartment, in contrast to *Bcl2l11*/BIM-/- phenotype^58^.

Expression of *Nr4as* and *Bcl2l11*/BIM are also induced during a second wave of negative selection triggered by TRA recognition in medullary SP thymocytes. BIM (along with PUMA) is essential for TRA-induced deletion in the medulla ^9,37,57,61^. We identify Nr4a consensus motifs and an Nr4a1/Nur77 ChIPseq peak in an enhancer required for *Bcl2l11* induction in the thymus whose deletion phenocopies *Bcl2l11*/BIM-/- and rescues negative selection ^62^. We also identify a profound competitive advantage for Nr4a DKO OT-II and polyclonal thymocytes at the SP stage that coincides with a complete failure of DKO OT-II thymocytes to upregulate *Bcl2l11*/BIM in the RIPmOVA model. In contrast, *Nr4a1* SKO OT-II thymocytes express subtly and non-significantly reduced levels of *Bcl2l11* transcript, suggesting functional redundancy between Nr4a1 and Nr4a3^16^. Taken together, our data are consistent with a model where strong TCR signaling in the medulla induces coordinate upregulation of Nr4a1 and Nr4a3, which trigger negative selection through transcriptional upregulation of *Bcl2l11*/BIM.

An extensive literature demonstrates that Treg fate is agonist-selected by expression of TRA in mTEC and can emerge at the boundary of positive and negative selection signaling thresholds^5^. CITEseq reveals that Treg and medullary SP thymocytes fated for negative selection occupy adjacent clusters with similar transcriptomes. For example, *Nr4a1, Nr4a3, Ikzf2*/HELIOS, and *Nt5e*/CD73 are expressed in both populations, suggesting they mark a common progenitor population of strongly “signaled” SP thymocytes. The fate decision between diversion to Treg and deletion is further influenced by additional signals, such as co-stimulation and IL-2^5^. Indeed, IL-2 supply has long been appreciated as a key and limiting requirement for Treg survival and fate in the thymus, and CITEseq analysis shows an IL2/STAT5 module marks this precursor population across a gradient. In contrast to a cell-intrinsic requirement for Nr4as to promote transition between CD25hi pre-Treg to CD25hi Foxp3+ Treg (evidenced by a profound block at this transition for DKO SP4 thymocytes)^64^, we observed a non-cell intrinsic impact of Nr4as on Treg fate. Rudensky and colleagues as well as others have suggested strongly-signaled thymocytes are the main source of IL-2 to promote Treg fate^45,47^; we postulate that DKO self-reactive clones that have escaped deletion are a source of excess IL-2 that promotes Treg fate among donor thymocytes of WT origin in the same chimeric thymus. In addition, Nr4a-deficient T cells have been shown by us and others to over-produce IL-2 ^19,66^.

Unexpectedly, in polyclonal DKO chimeras we describe emergence of a unique Foxp3+ SP8 population (as well as SP8 CD25hi Foxp3-precursors) that are absent in WT chimeras. Foxp3+ SP8 is both a cell-intrinsic and cell non-intrinsic phenotype in DKO chimeras, raising the possibility that both escape of DKO SP8 from deletion and excessive IL-2 production may contribute to this fate. This may correspond to a Foxp3+ CD8+ peripheral population identified among tumor-infiltrating T cells as well as in alloreactive T cells from GVHD models where it is constrained by high BIM expression^48,49^. Indeed, we show that semimature SP8 thymocytes express higher levels of both Nur77-GFP and BIM, and are more stringently censored by deletion than SP4 thymocytes^43,44^. Strikingly, BIM expression is suppressed in both a cell-intrinsic and cell non-intrinsic manner in SP8 thymocytes from polyclonal DKO chimeras. Nr4as are required to induce *Bcl2l11* transcript, but excess IL-2 produced by DKO thymocytes may suppress *Bcl2l11* in WT cells; indeed IL-2 has been reported to prevent deletion of Foxp3+ T cells, and their survival in Il2-/- animals is rescued by *Bcl2l11*-/-^46,82^. Taken together, this leads us to propose that CD8 Treg fate is latent but normally censored by deletion in the thymus and that both Nr4a and IL-2 signals may converge on BIM to do so.

Importantly, in RIPmOVA hosts, WT OT-II thymocytes are fated for deletion rather than Treg diversion; the overwhelming majority of Treg emerge from polyclonal and not OT-II donor populations in competitive chimeras. Our transcriptomic analyses indicate that neither *Il2ra*/CD25 nor *Foxp3* are substantially upregulated in OT-II cells, and therefore the strongly signaled SP cells likely do not comprise a population of committed pre-Treg - in contrast to prior studies ^64^. Rather, we postulate that HELIOS and CD73 expression mark all strongly signaled DKO thymocytes that have escaped deletion. A similar HELIOS+ population was identified in *Bcl2l11*/BIM-/- InsHEL model even after Treg are gated out ^37^. Strikingly, DKO naive CD4 T cells in the periphery with a similar phenotype (CD73+HELIOS+) emerge at the same very early time points. These cells exhibit not only phenotypic but also functional features of anergy yet are harvested from RIPmOVA host spleens. The data lead us to postulate that this ‘naïve’ peripheral T cell anergic phenotype is induced by thymic rather than peripheral self-antigen encounter.

We discovered that the Nr4a family is an essential molecular link between TCR signaling and a broad transcriptional program induced in response to TRA. Interestingly, a thymic transcriptional defect reminiscent of DKO thymocytes has been described in autoimmune-prone NOD mice. NOD thymocytes exhibit impaired *Bcl2l11* induction and escape from deletion triggered by InsHEL model antigen^35,60^. Of the 82 InsHEL-induced genes that overlap with RIPmOVA-induced genes in our data set, over half (56) exhibit a defect in the NOD background. Since this defect is broad but not global, NOD – like DKO thymocytes – does not reflect a simple proximal defect in TCR signal transduction. Among NOD-dependent genes, we observed a remarkable enrichment in Nr4a-dependent genes (23/56), including *Eno3, Tnfrsf9, Lad1*, and of course *Bcl2l11* itself. We propose that polygenic defects affecting the Nr4a-dependent thymic transcriptome may confer risk for autoimmunity.

The broad transcriptional program induced by RIPmOVA TRA in thymocytes overlaps with TCR-induced programs in mature T cells. However, only a small number of induced transcripts correspond to primary response genes. Rather, the majority require chronic antigen stimulation for their expression in the periphery and are associated with anergy and Treg fate. This “anergy” signature includes genes upregulated in exhausted CD8 T cells and tolerant CD4 T cells (signal 1 without signal 2), as well as NFAT-induced genes. Notably, Nr4as have been shown to contribute to the anergy/exhaustion transcriptome and epigenome of peripheral T cells^19,66,83^ and here we show that they may contribute to a related transcriptional program of non-deletional tolerance in the thymus as well.

Functional unresponsiveness as an alternative fate to deletion in the thymus has been previously proposed; classic papers from over three decades ago showed this for class I-restricted TCRs ^84^ or class II-restricted TCRs ^85^ that escape deletion by super-antigen (Mls-1a), and also for TEC expression of an alloantigen driven by keratin IV promoter ^86^. More recently a tetramer approach was used to demonstrate a variety of self-reactive T cell fates (spanning deletion, diversion, and ‘ignorance’) driven by varying location and amount of self-antigen presentation in the thymus ^87^. Elimination of negative selection has also unmasked alternate fates for self-reactive lymphocytes^88^. *Bcl2l11*/BIM-deficient T cells, which escape deletion by TECs yet do not produce autoimmune disease ^57,61^, express anergic phenotypic markers CD73/FR4 in the periphery ^58^, and exhibit impaired TCR signaling^89^. However, it has not been clear in most prior studies if clonal anergy of cells evading deletion was imposed in the thymus or only in the periphery where it has long been appreciated to play a vital role. Most recently, Baldwin and colleagues showed that non-deletional control of OTI *Bcl2l11*/BIM-/- thymocytes from RIPmOVA hosts is associated with PD-1 upregulation in the thymus^90^. We propose that *Pdcd1*/PD1 induction in response to self-antigen is part of a larger transcriptional program we identify in RIPmOVA OT-II thymocytes that is partly mediated by the Nr4a family. Our DKO chimera model uniquely disables both Treg diversion and deletion to unmask this tolerance program in the thymus and the contribution of the Nr4as to its establishment.

Naturally occurring naïve (CD62Lhi) peripheral T cells marked by high Nur77-eGFP expression exhibit impaired TCR signaling, IL-2 production, and an increased propensity to differentiate into Treg ^71,72^. Strikingly, we show that this population is enriched for recent thymic emigrants and also harbors an epigenetic imprint that corresponds to the Ag-induced and Nr4a-dependent transcriptional program we identified in self-reactive SP thymocytes. Expression of inhibitory co-receptors such as Pdcd1/PD1, Ctla4, Lag3, and Cd200 in this thymically-imprinted RTE population might render these cells particularly vulnerable to checkpoint blockade. Consistent with this idea, it was recently shown that thymoma and thymic remnant size constituted major factors for development of checkpoint myocarditis and myositis ^91^. The anergy-like transcriptional program we describe in self-reactive thymocytes may represent a poised state that operates to preserve tolerance at the boundary of Treg diversion in the thymus and can persist among RTEs after thymic egress– enabling clones that encounter peripheral self-antigen to undergo diversion, deletion, and anergy^92^. Anergic CD73hiFR4hi T cells in the CD44hiCD62Llo memory compartment are also poised to differentiate into Treg suggesting a transcriptional and epigenetic link between anergy and Treg persists throughout T cell ontogeny^70^.

By transmitting a broad TCR-induced transcriptional program in response to TRA recognition in the medulla, Nr4as not only mediate Treg diversion as previously shown, but also drive BIM-dependent deletion, and contribute to a non-deletional anergy-like tolerance program in the thymus that can be identified in the periphery.

## Materials and methods

### Mice

Mice were housed in a specific pathogen-free facility at The University of California in San Francisco according to University and NIH guidelines. C57BL/6 (CD45.2) and BoyJ (CD45.1) mice (both considered WT here) were initially purchased from The Jackson Laboratory or Charles River Labs, respectively. Nur77-eGFP BAC Tg mice were previously described^24^. OT-II TCR transgenic mice^93^ and RIP.mOVA transgenic mice^50^ were previously described. *Nr4a3*-deficient allele was generated in our laboratory on the C57BL/6 genetic background as previously described ^75^ *Nr4a3*-/-*Nr4a1*-/- (DKO) mice were previously described^19^ and were bred to OT-II mice to create the OT-II *Nr4a1*-/-*Nr4a3*-/- strain. BIM-deficient B6.129S1-*Bcl2l11^tm1.1Ast^*/J line (Jackson #:004525)^9^ were crossed to Nur77-GFP mice. Foxp3-RFP^94^ mice crossed to Nur77-GFP were previously described^72^. All strains were fully backcrossed to C57BL/6 genetic background for at least 6 generations. RIP.mOVA CD45.1/2 and WT CD45.1/2 mice were F1 progeny from RIP.mOVA x BoyJ crosses. OTII was crossed with BoyJ to generate and maintain OTII CD45.1 line. Mice of both sexes were used for experiments between the ages of 3 and 10 weeks except for BM chimeras as described below. Study approval. All mice were housed in a specific pathogen-free facility at UCSF and Emory University according to the University and National Institutes of Health guidelines.

### Antibodies and Reagents

Abs for surface markers: Abs to CD3, CD4, CD5, CD8, CD19, CD24, CD25, CD44, CD45.1, CD45.2, CD62L, CD69, CD73, FR4, Vα2, Vβ5, and TCRβ conjugated to fluorophores were used (BioLegend, eBiosciences, BD, or Tonbo). Abs for intra-cellular staining: HELIOS Ab conjugated to PE (clone 22F6, cat. 563801, BD Biosceinces), BIM (clone C34C5; cat. 2933S, Cell Signaling) Rabbit mAb, FOXP3 Ab conjugated to APC or FITC (clone FJK-16s, Invitrogen). Anti-pERK (Phospho-p44/42 MAPK (T202/Y204) (clone 197G2, Cell Signaling) Rabbit Ab. Donkey Anti-Rabbit IgG (H+L) conjugated to APC (Jackson ImmunoResearch). Stimulatory Abs: Anti-CD3 (clone 2c11) (BioLegend). Goat anti-Armenian Hamster antibody (Jackson ImmunoResearch).

Culture Media: RPMI-1640 + L-glutamine (Corning-Gibco), Penicillin Streptomycin L-glutamine (Life Technologies), HEPES buffer [10mM] (Life Technologies), B-Mercaptoethanol [55mM] (Gibco), Sodium Pyruvate [1mM] (Life Technologies), Non-essential Amino acids (Life Technologies), 10% heat inactivated FBS (Omega Scientific).

### Flow Cytometry

Cells were analyzed on a Fortessa and sorted on Aria (Becton Dickson). Data analysis was performed using FlowJo (v9.9.6 or v10.7.1) software (Becton Dickson).

### FOXP3/HELIOS staining

FOXP3 and HELIOS staining was performed utilizing a FOXP3/transcription factor buffer set (eBioscience) in conjunction with APC or FITC anti-FOXP3, as per manufacturer’s instructions.

### BIM staining

Intracellular staining for BIM was performed with secondary Donkey anti-rabbit APC secondary antibody using BD cytofix/cytoperm kit (BD biosciences 554722) per manufacturer’s instructions.

### Live/dead staining

LIVE/DEAD Fixable Near-IR Dead Cell Stain kit (Invitrogen). Reagent was reconstituted in DMSO as per manufacturer’s instructions, diluted 1:1000 in PBS, and cells were stained at a concentration of 1 × 106 cells /100 μl on ice for 15 minutes.

### Bone marrow chimeras

Chimeras were generated by irradiating hosts either with Cesium source (2 doses of 530 cGy, 4 hours apart) and injection on the same day IV with a total of 2 × 10^6^ donor BM cells or with an X-ray source (2 doses of 450 cGY, 3-4 hours apart) and rescued with IV transfer of 5 × 10^6^ donor BM cells the following day. For each chimeras type, BM from two or three congenically marked donor strains was mixed at specified ratios for transfer as depicted in schematics (Figs 1c, 2a, 3a). For polyclonal chimeras generated with X-ray source (Fig 1c design for experiments depicted in Figs 1, 8, 9), both CD45.2 DKO and WT polyclonal donor marrow was depleted of mature T cells by magnetic column-based negative selection (Miltenyi Biotec MACS) using an anti-mouse CD5 biotin-conjugated antibody (Clone 53-7.3, Miltenyi Biotec, 130-101-960) and Streptavidin MicroBeads (Miltenyi BioTec, 130-048-102) per manufacturer.s instructions. Chimeras were assessed between 5-11 weeks post-reconstitution as noted in figures except for Fig 9h, i phosflow assays which were conducted with polyclonal chimeras at 14 weeks.

### Phospho-flow

Splenocytes were rested at 37°C in serum-free RPMI for 30 minutes. Cells were then stimulated with 10 μg/ml of anti-CD3 (clone 2c11) for 30 seconds followed by 50 μg/ml of anti-Armenian hamster crosslinking antibody for 2 minutes, or PMA for 2 minutes. Stimulated cells were fixed with 2% paraformaldehyde and permeabilized with methanol at -20°C overnight. Cells were then stained with surface markers and pErk at 20°C.

### Histology

Organs were harvested into 10% formalin for 24hrs, dehydrated progressively, stored in ethanol at 4C until processing for H&E by Histowiz.

### Data visualization, analysis, and statistics

Flow cytometry data was analyzed using FlowJo software FlowJo v10.0 Software (BD Life Sciences). Graphs and statistical analyses were undertaken using Prism v6. For RNAseq data, see below.

### RNA sequencing

DKO OT-II:OT-II:WT -> WT hosts and DKO OT-II:OT-II:WT -> RIP.mOVA hosts bone marrow chimeras were created as described above (Fig 3a schematic). DKO OT-II (CD4+ CD8-CD69+ Vα2+ CD45.1-CD45.2+) and control OT-II (CD4+ CD8-CD69+ Vα2+ CD45.1+ CD45.2+) semimature SP thymocytes were isolated on BD FACSAria II and BD FACSAria Fusion cell sorters directly into RLT + 1% beta-mercaptoethanol (BME) buffer (Qiagen). Libraries were generated by the Emory Integrated Genomics Core (EIGC) as follows: RNA was isolated using the Quick-RNA MicroPrep kit (Zymo,11-328M). 2000 cell equivalent of RNA was used as input to SMART-seq v4 Ultra Low Input cDNA Synthesis kit (Takara, 634888) and 200 pg of cDNA was used to generate sequencing libraries with the NexteraXT kit (Illumina, FC-121-10300). Libraries were quantified by qPCR and bioanalyzer traces, pooled at equimolar ratios, and sequenced on the NovaSeq6000 with a PE100 configuration using a NovaSeq 6000 SP Reagent Kit. Raw fastq files were mapped to the mm10 genome using STAR^95^ with the GENCODE vM17 reference transcriptome. PCR duplicate reads were marked with Picard MarkDuplicates and removed from downstream analyses. Reads mapping to exons for all unique ENTREZ genes were summarized using GenomicRanges^96^ in R v3.5.2 and normalized to reads per kilobase per million. Differentially expressed genes were determined using DESeq2^97^, and genes that displayed an absolute log2 fold change >1 and Benjamini–Hochberg false discovery rate (FDR)–corrected *p* value <0.05 were considered significant.Heatmaps were generated using the Morpheus online tool from Broad (https://software.broadinstitute.org/morpheus/) and volcano/FC plots were generated using R Statistical Software. Fastq and RPKM data are publicly available and can be accessed from NCBI’s Gene Expression Omnibus using the GEO Series access number GSE235101.

## Acknowledgments

We thank Wan-Lin Lo, Ellen Robey, and Arthur Weiss for helpful discussions and critical reading of the manuscript. We thank Alfonse Roque for help with mouse husbandry.

---

**Supp Figure 1.**
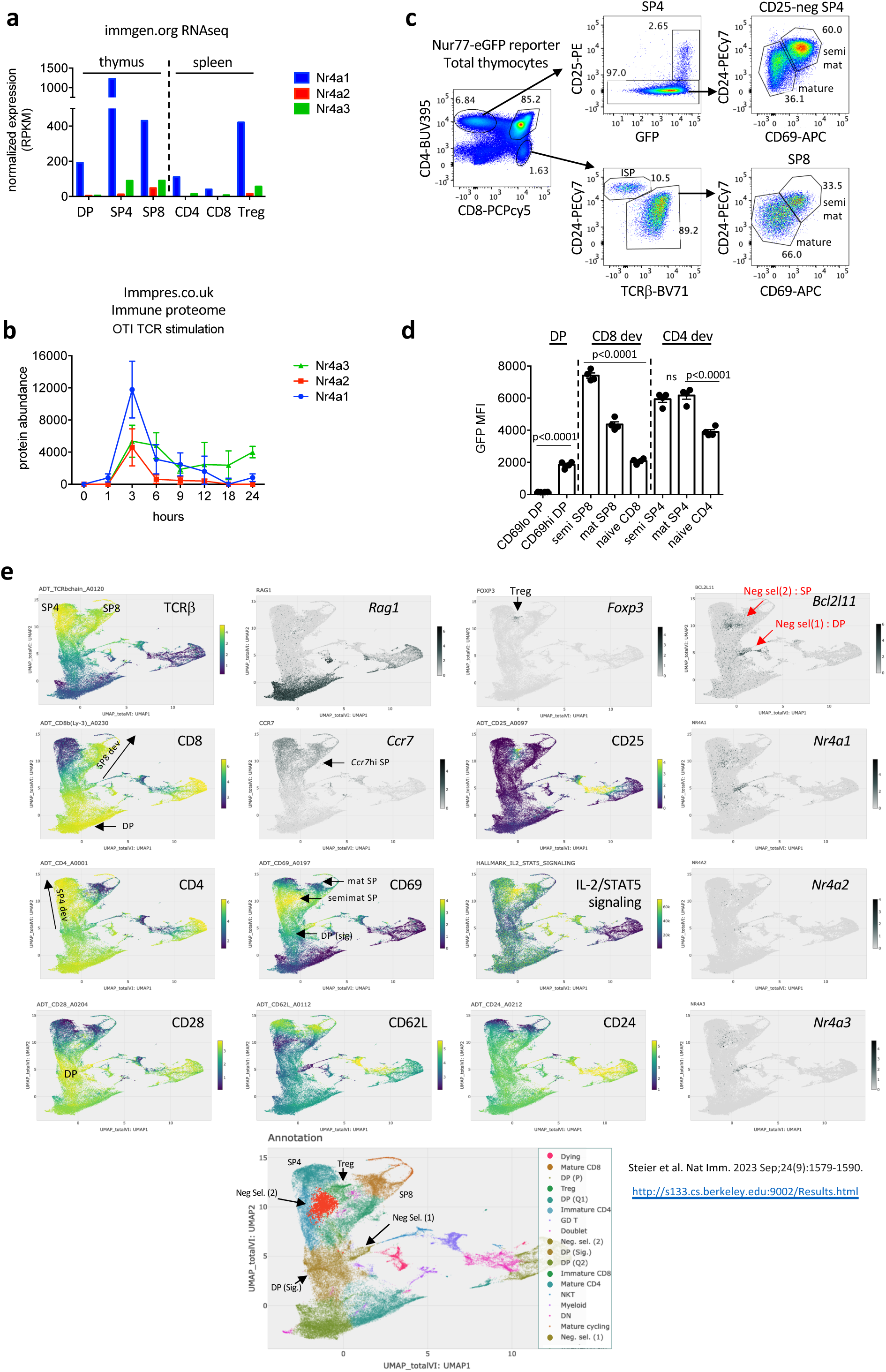
**Related to Main Figure 1** **a.** RPKM from public RNAseq data sets are plotted (immgen.org) depicting Nr4a1, 2, 3 transcript abundance among T cell lineage subsets in thymus and periphery. **b.** mean +/- SD values from immune proteome database (Immpres.co.uk) are plotted depicting mass spec abundance of Nr4a1/2/3 protein products in TCR stimulated CD8 T cells across a time course. **c.** Gating scheme depicts identification of DP and SP subsets among Nur77-eGFP reporter thymocytes. **d.** Quantification of GFP MFI among Nur77-eGFP reporter thymic subsets as gated in c, in 4 biological replicates +/- SEM **e.** Selected protein and transcript UMAP data from publicly available VISION thymus CITEseq interface (http://s133.cs.berkeley.edu:9002/Results.html; Steier et al. Nat Imm. 2023 Sep;24(9):1579-1590.) <u>Statistical tests:</u> d. one way ANOVA with Holm-Sidak correction for multiple pre-selected comparisons. No matching or pairing. Data in c, d representative of at least 4 biological replicates

**Supp Figure 2.**
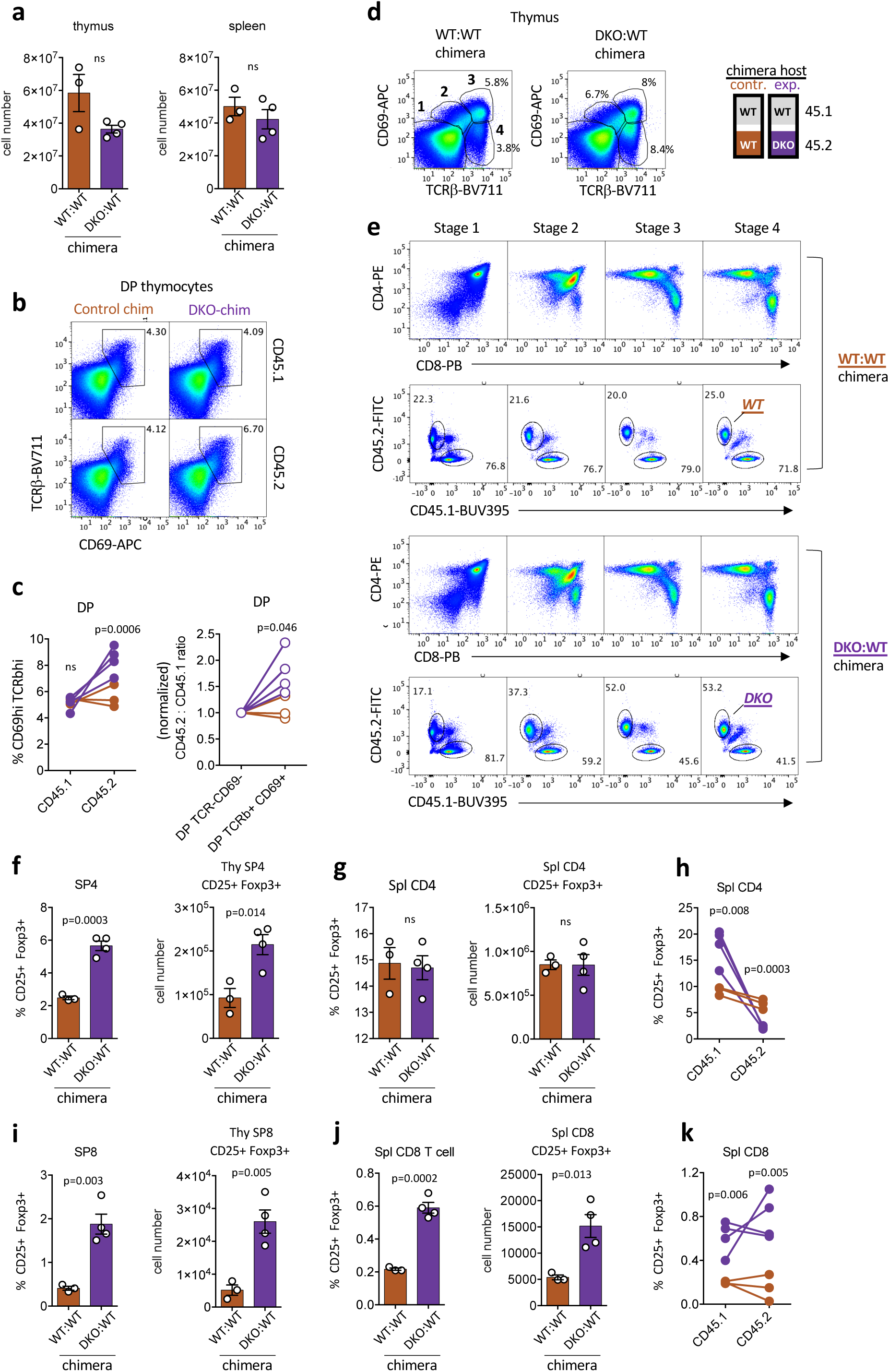
**Related to Main Figure 1** **a.** Graphs depict absolute cell number in thymus or spleen of polyclonal chimeras (see schematic Fig 1c) in biological replicates +/- SEM **b, c.** Representative plots depict DP thymocytes of each donor genotype in polyclonal control and experimental chimeras (schematic in Fig 1c) stained to identify pre- and post-selection gates defined by CD69 and TCRβ expression. Graphs show quantification of gates (left) or ratio of donor genotypes (normalized to pre-selection DP) among these gates for biological replicates. **d, e.** Representative plots (e) depict co-receptor agnostic gating to identify sequential stages of thymocyte development 1-4. In (e) thymic stages 1-4 as gated in (d) are shown for WT:WT (top) and DKO:WT (bottom) polyclonal chimeras (see schematic 1c). Both coreceptor expression (top) and CD45.1/CD45.2 donor genotypes (bottom) are depicted for each stage. **f, g.** Graphs depict total % and absolute number of Foxp3+ CD4+ Treg (irrespective of donor genotype) from thymus and spleen of experimental or control polyclonal chimeras (schematic in Fig 1c) in biological replicates +/- SEM. **h.** Graph depicts % Foxp3+ CD4+ Treg of each donor genotype in polyclonal chimeras. Lines depict donors from the same chimera. **i, j.** Graphs depict total % and absolute number of Foxp3+ CD8+ cells (irrespective of donor genotype) from thymus and spleen of experimental or control polyclonal chimeras (schematic in Fig 1c) in biological replicates +/- SEM. **k.** Graph depicts % Foxp3+ CD8+ cells of each donor genotype in polyclonal chimeras. Lines depict donors from the same chimera. <u>Statistical tests</u>: a, d, e, f, g, unpaired two-tailed paramtetric t-test, Assume equal SD c, h, i unpaired two-tailed t-test, do not assume equal SD Graphs in this fig depict N=3 control chimeras and N=4 polyclonal chimeras and are representative of 3 independent sets of chimeras.

**Supp Figure 3.**
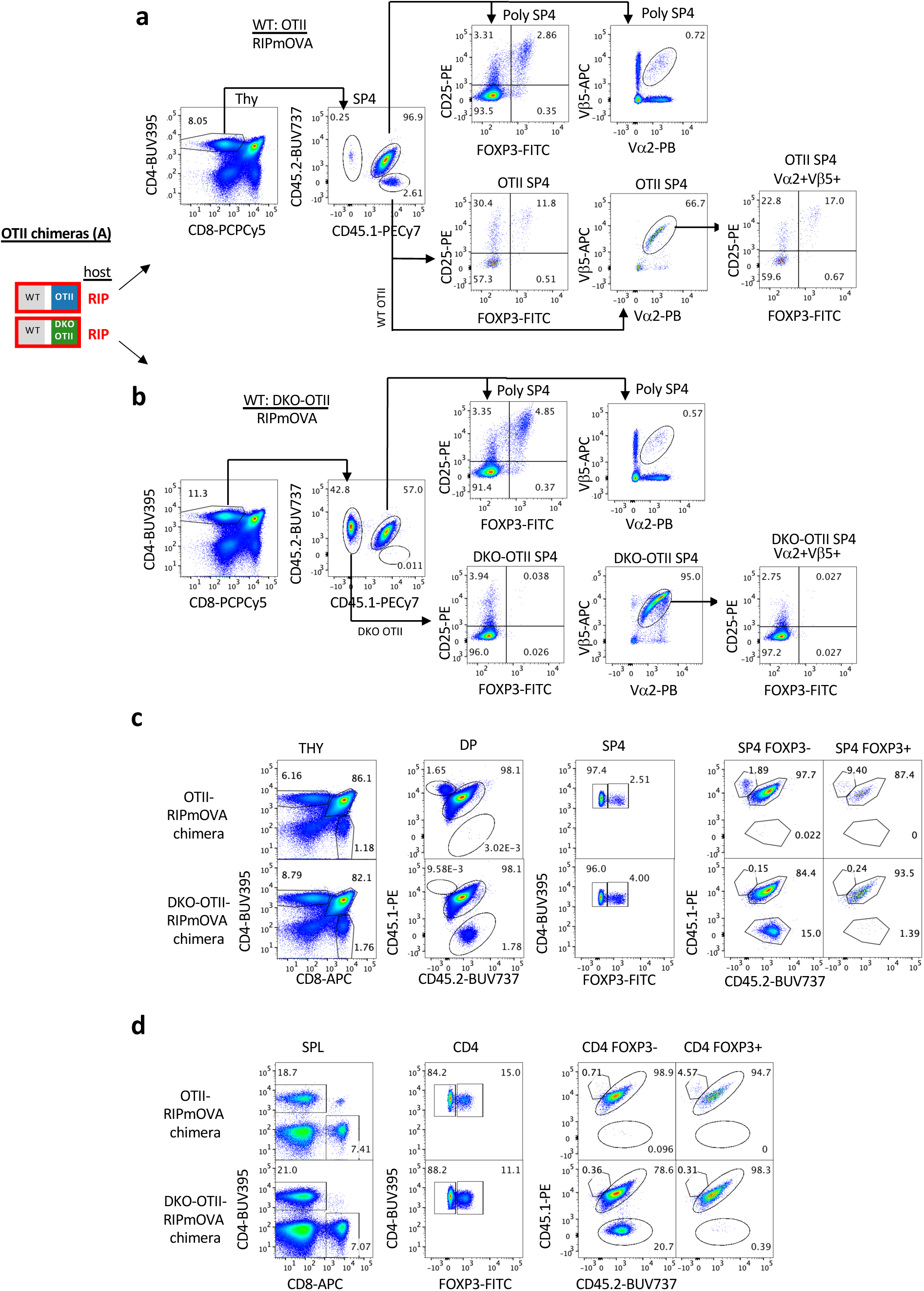
**Gating scheme related to Main Figure 2** a-d. Gating schemes corresponding to chimera schematic in Main Fig 1b. Gating schemes in (a, b) depict donor genotypes (WT-OTII or DKO-OTII as well as admixed polyclonal donor) within SP4 compartment, and arrows direct to CD25/Foxp3 staining for each genotype to define Treg, pre-Treg, Tconv gates followed by Va2 and Vb5 staining to exclude escapees. Gating in (c, d) depict inverted gating scheme depicting contribution of donor genotypes in chimeras to each gate DP, SP4 Foxp3-, and SP4 Foxp3+ Treg.

**Supp Figure 4.**
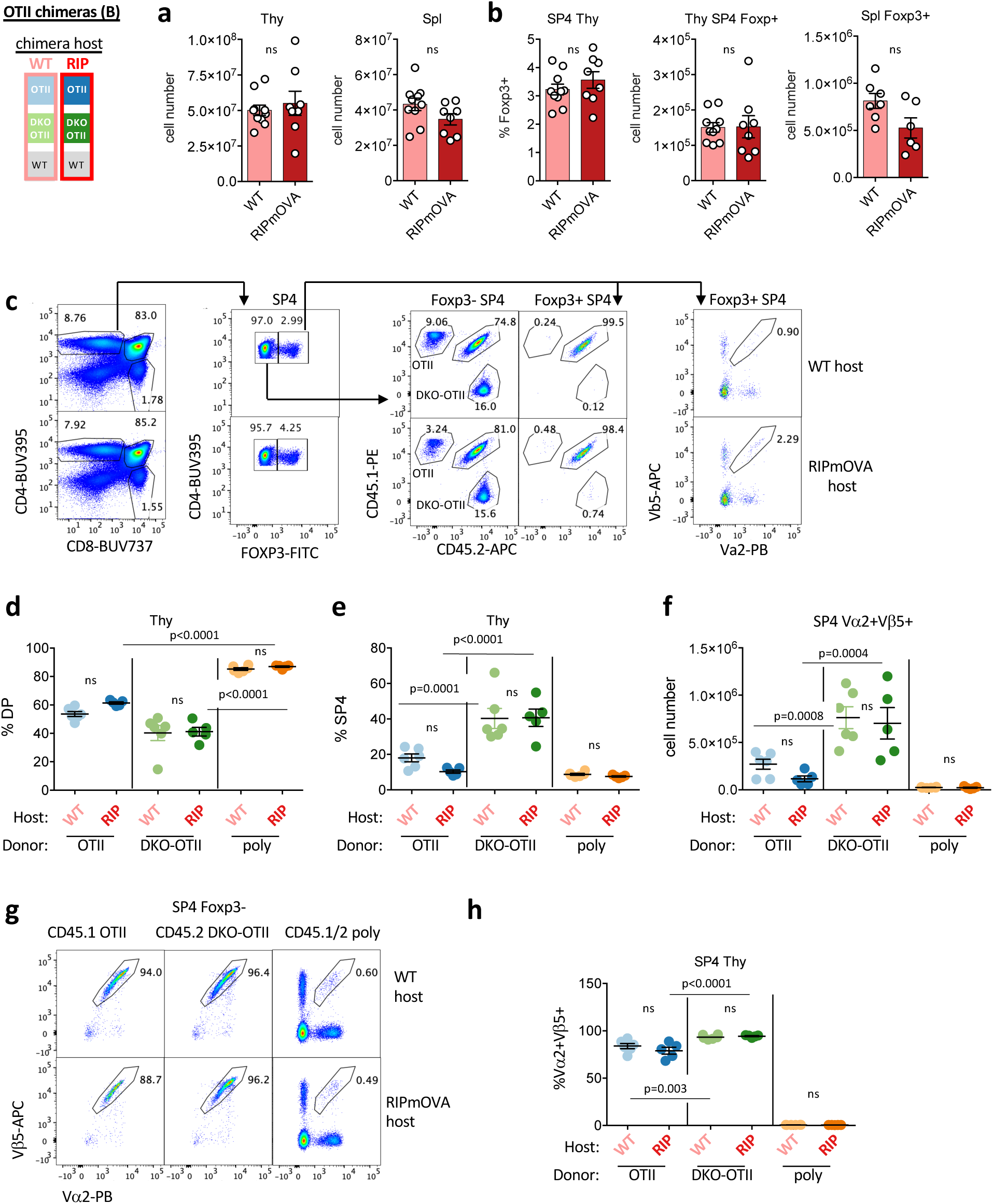
**Related to Main Figure 3** **a.** Graphs depict absolute cell number in thymus or spleen of RIPmOVA and WT host chimeras (see schematic Fig 3a) in biological replicates +/- SEM **b.** Graphs depict relative and absolute number of Foxp3+ Treg in thymus and periphery of RIPmOVA and WT host chimeras (see schematic Fig 3a) in biological replicates +/- SEM **c.** Representative gating scheme depicts donor genotypes among Foxp3- and Foxp3+ SP4 thymocytes as well as surface TCR chain expression among Foxp3+ SP4. **d, e.** Graphs depict quantification of DP and SP4 populations from donor genotypes from WT host and RIPmOVA host chimeras (see Fig 3a schematic) as gated in Fig 3 for biological replicates +/- SEM. **f.** Graphs depicts absolute number of Vα2+Vβ5+ SP4 populations from donor genotypes from WT host and RIPmOVA host chimeras (see Fig 3a schematic) **g, h.** Representative plots depict Foxp3-SP4 thymocytes of each donor genotype from WT host and RIPmOVA host chimeras (see Fig 3a schematic) stained and gated to detect Vα2+Vβ5+ cells. Graph (h) depicts quantification of gating in (g) for biological replicates +/- SEM. <u>Statistical tests:</u> a, b, Unpaired 2-tailed parametric t test, assume equal SD. d, one way ANOVA with Tukey’s multiple hypothesis test. e, f, h, one way ANOVA with pre-specified comparisons corrected for by Sidak test

**Supp Figure 5.**
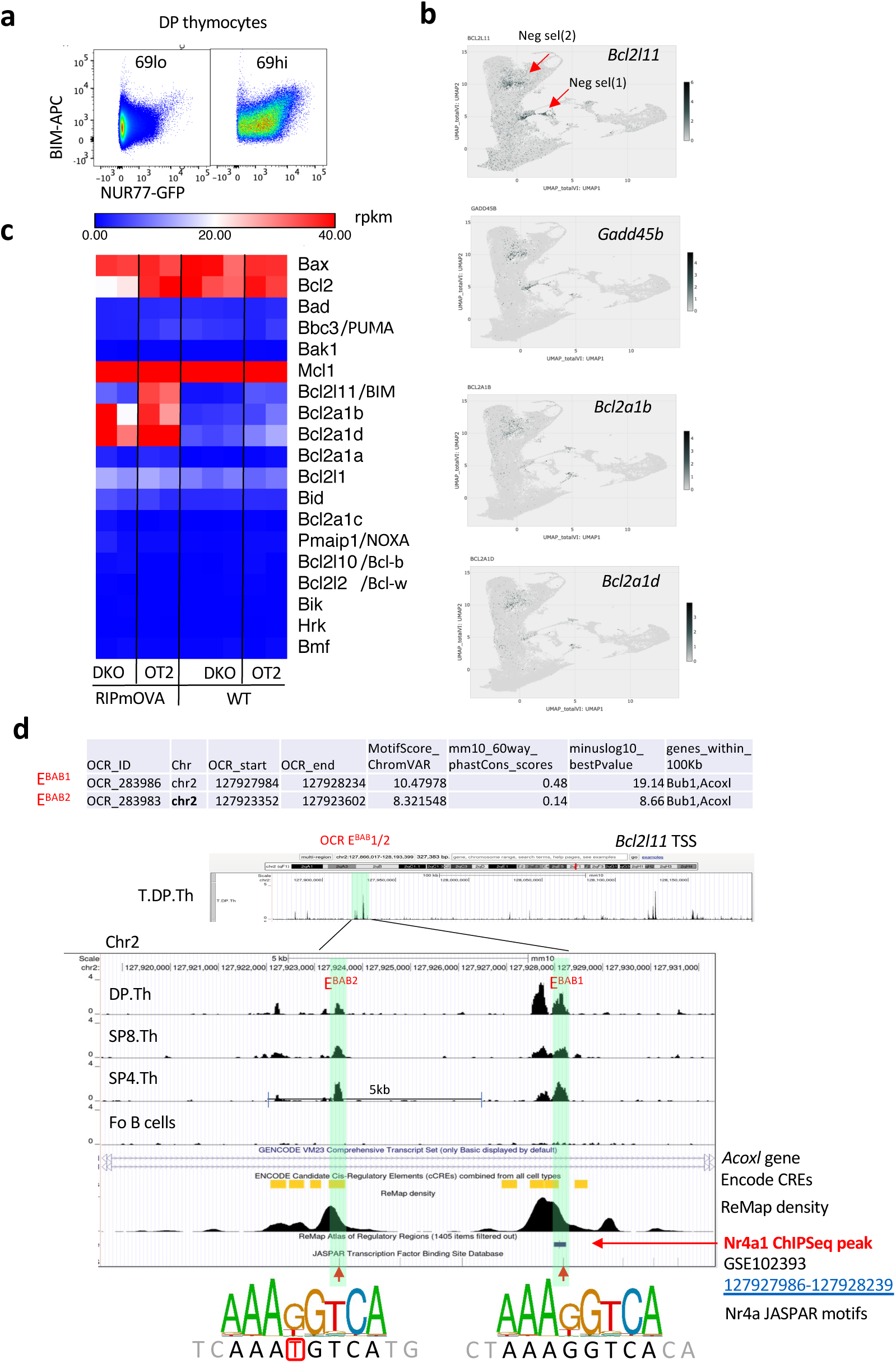
**Related to Main Figure 4** **a.** DP thymocytes as described in Fig 4b are gated to identify pre-selection CD69lo and post-selection CD69hi subsets. Representative plots depict BIM and Nur77-GFP expression. **b.** Selected transcript UMAP data from publicly available VISION thymus CITEseq interface (http://s133.cs.berkeley.edu:9002/Results.html; Steier et al. Nat Imm. 2023 Sep;24(9):1579-1590.) **c.** Heatmap depicts absolute rpkm values for pro- and anti-apoptotic Bcl2 family members from RNAseq data set schematized in Fig 3a and described in Fig 5a (Table 1a) **d.** Genomic coordinates of E^BAB1^ and E^BAB2^ OCRs; UCSC browser tracks depict ATACseq peaks from public immgen.org as shown in Fig 4i but zoomed in.

**Supp Figure 6.**
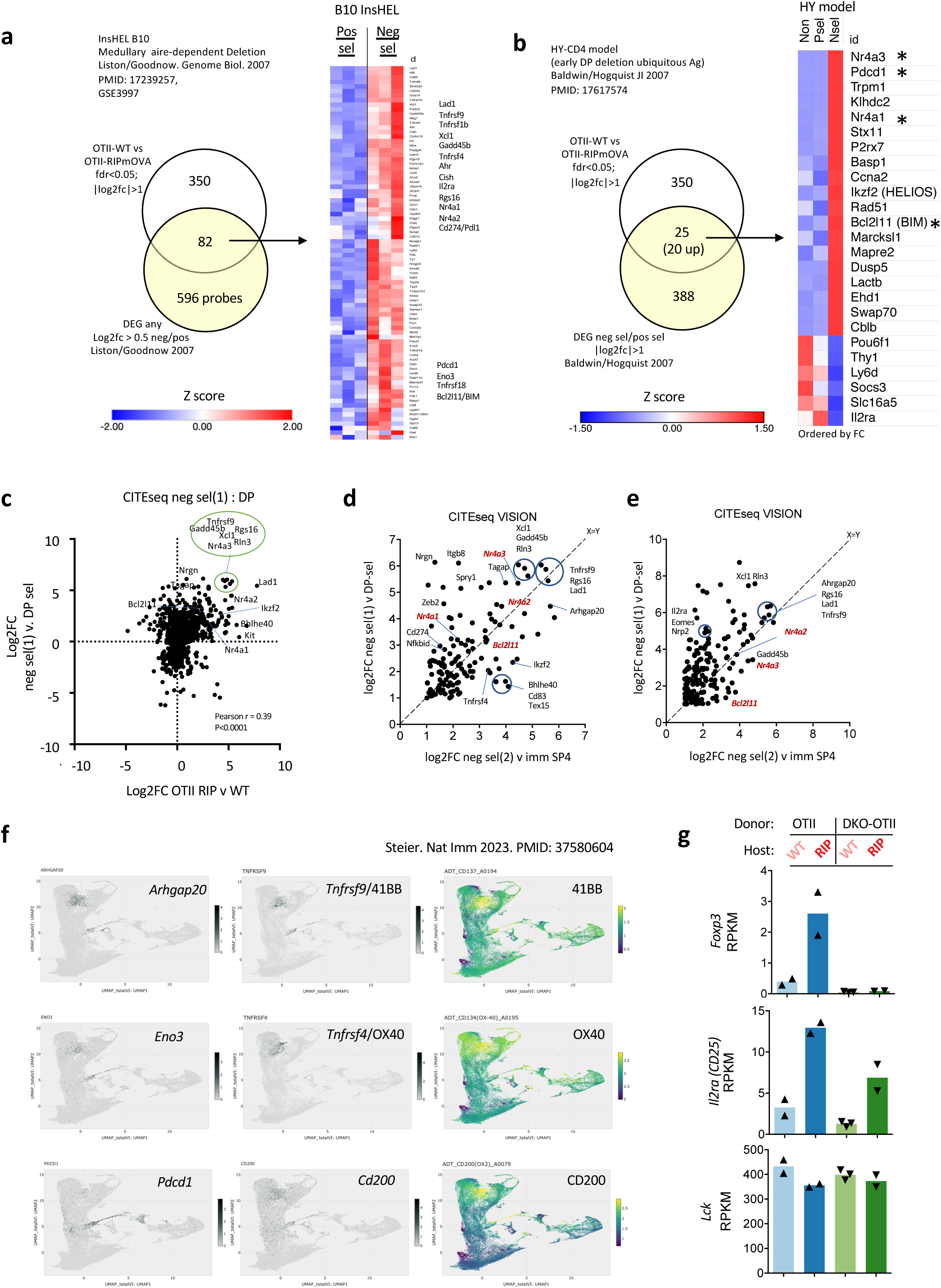
**Related to Main Figure 5** **a.** Venn diagram depicts overlap between OTII-RIPmOVA DEG (N=350 as in Fig 5a) and InsHEL model (PMID: 17239257, GSE3997). Heatmap depicts overlapping gene expression in InsHEL model. see also Table 2a. **b.** Venn diagram depicts overlap between OTII-RIPmOVA DEG as above and HY-CD4 model (PMID: 17617574). Heatmap depicts overlapping gene expression in HY-CD4 model. see also Table 2b **c.** FC FC plot as in Fig 5b except CITEseq comparator depict Log2FC Neg sel (1) cluster / signaled DP. Pearson correlation coefficient is quantified. See also Table 2c. **d, e.** FC FC plots as in c above except comparing Neg Sel clusters 1, 2 to non-deleting closest comparitors (signaled DP and imm SP4 respectively). Genes upregulated in Neg Sel and Treg are included in the plot. See also Table 2d. **f.** Selected protein and transcript UMAP data from publicly available VISION thymus CITEseq interface (http://s133.cs.berkeley.edu:9002/Results.html; Steier et al. Nat Imm. 2023 Sep;24(9):1579-1590.) **g.** Graphs depict RPKM for *Foxp3, Il2ra*, and *Lck* in CD69hi Va2+ SP4 thymocytes of either WT-OTII or DKO-OTII donor genotypes from chimeras depicted in 3a. RNAseq was performed on biological replicates (Table 1a depicts p value and fdr for all pairwise comparisons across sample types, GSE235101).

**Supp Figure 7.**
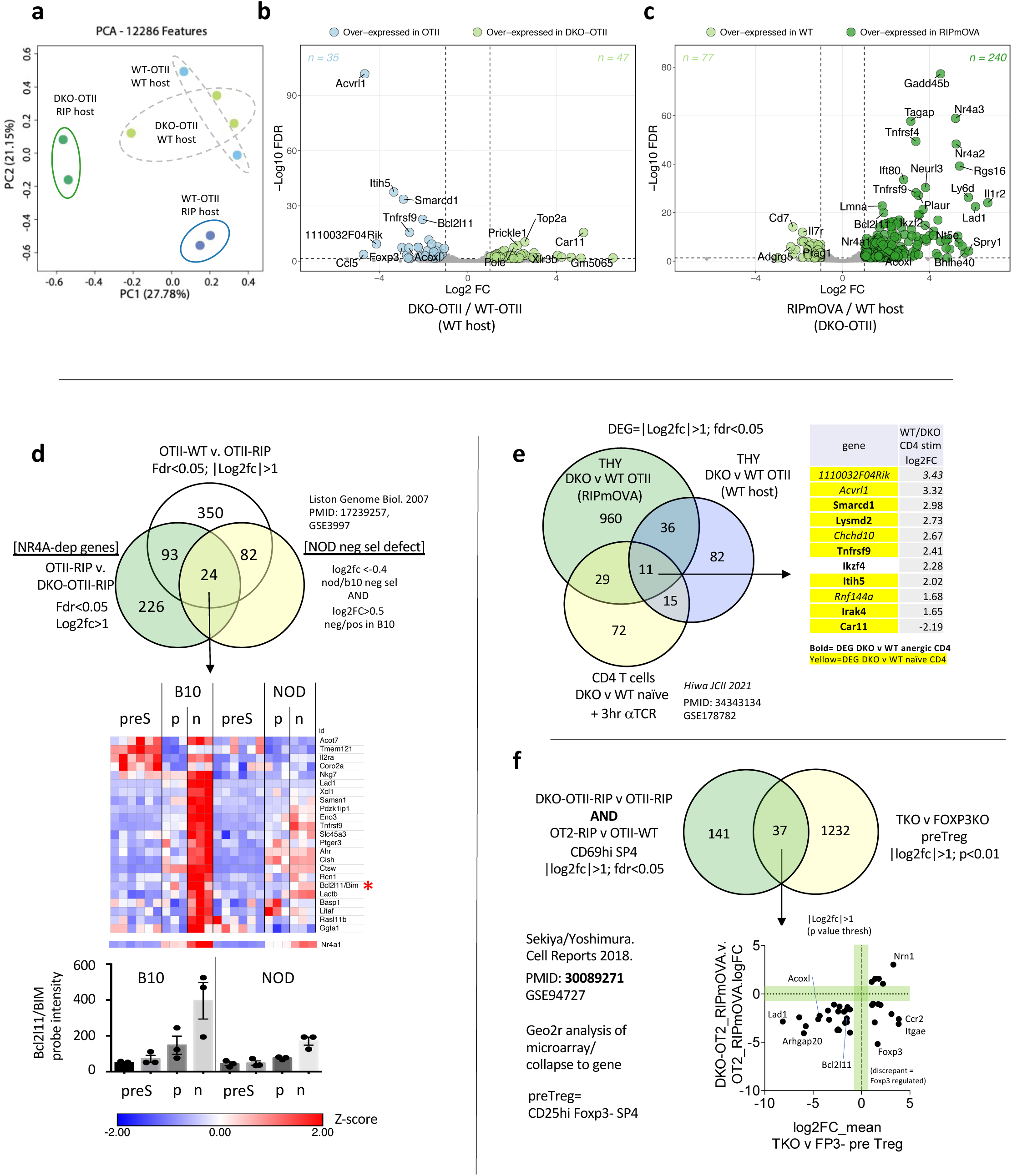
**Related to Main Figure 5** **a.** PCA plot depicts data from RNAseq experiment described in Fig 5a, GSE235101) **b.** As in (a) except volcano plot depicts DEG (colored points) in WT-OTII vs. DKO-OTII thymocytes sorted from WT hosts. **c.** As in (a) except volcano plot depicts DEG (colored points) DKO-OTII thymocytes sorted from either WT or RIPmOVA hosts. **d.** Venn diagrams depict overlap between Nr4a-dependent genes and those induced by RIPmOVA in our RNAseq data set (white/green circle overlap) along with NOD-dependent genes induced in InsHEL model (yellow circle; PMID: 17239257, GSE3997). Filtering criteria as noted in figure. See also Table 2a **e.** Venn diagrams depict Nr4a-dependent genes in thymocytes from RIPmOVA and WT host chimeras (white, blue circles) compared to Nr4a-dependent genes from peripheral naïve (CD62Lhi CD25- CD73- FR4-) CD4 T cells (DKO v WT) stimulated through TCR for 3 hr (yellow circle, PMID: 34343134; GSE178782). Filtering criteria as noted in figure: =|Log2fc|>1; fdr<0.05. see also Table 2g which additionally identifies shared Nr4a-dependent genes among naïve CD4 ex vivo and among anergic CD4 (CD44hi CD62Llo CD25-CD73hi FR4hi) of same data set. Overlap among all data sets annotated with bold, yellow highlighting. **f.** Venn diagram depicts Nr4a-regulated genes induced in RIPmOVA hosts in our data set (white circle) with Nr4a-dependent genes in pre-Treg (CD25hi Foxp3- thymocytes) defined by compqring Nr4ra1/2/3 TKO with Foxp3KO (PMID: 30089271; GSE94727 microarray data set analyzed via Geo2r and collapsed to gene). Filtering criteria as noted in figure. FC FC plot depicts overlap genes to identify concordantly Nr4a-dependent genes in bottom left quadrant. See also Table 2f

**Supp Figure 8.**
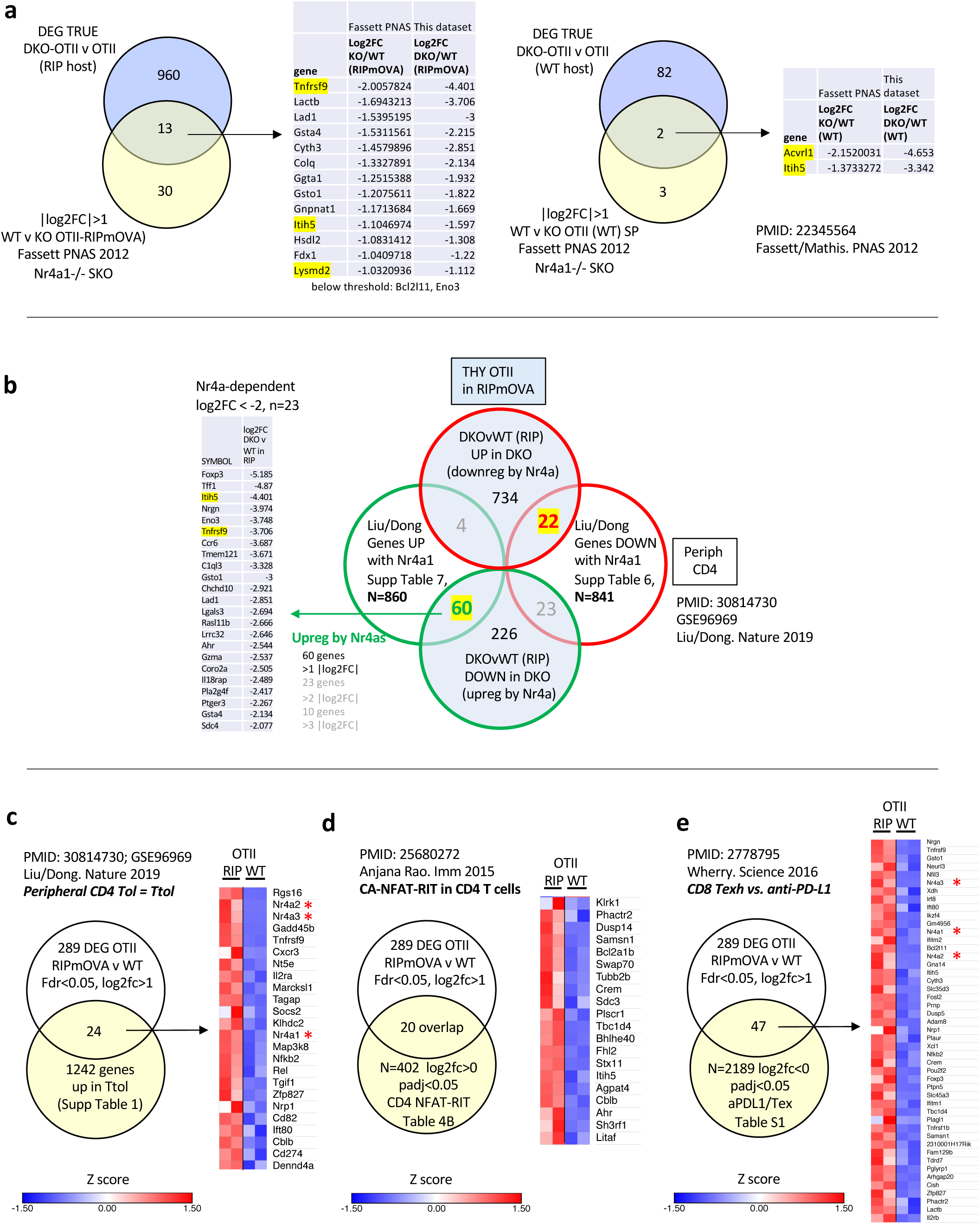
**Related to Main Figure 6** **a.** Venn diagrams compare DKO-dependent genes in either RIPmOVA (left) or WT (right) host chimeras with analogous samples from Nr4a1-/- OTII expressing RIPmOVA or not. Filtering criteria as noted in figure. Log2FC of overlap genes from both data sets as noted. **b.** Venn diagrams compare DEG downregulated (top red/blue circle) or upregulated (bottom green/blue circle) between DKO-OTII and WT-OTII thymocytes from RIPmOVA host chimeras against Nr4a-dependent genes identified in peripheral T cells using Nr4a misexpression (white circles with red/green outlines; PMID: 30814730; GSE96969; filtering as applied by authors to generate Tables 6, 7 in reference). Yellow highlighting identifies concordantly Nr4a-regulated genes across data sets and table shows most highly Nr4a-upregulated shared genes. see also Table 2h **c.** Venn diagram depicts overlap between RIPmOVA-induced genes in our data set (white circle, filtering criteria as noted in figure) and Ttol genes (upregulated in vitro in CD4 T cells by APC lacking B7 ligands, blue circle, PMID: 30814730; GSE96969; filtering as applied by authors to generate Table 1 in reference). Heatmap depicts expression of overlap genes in our data set and asterisk highlights *Nr4a1/2/3*. see also Table 2j **d.** Venn diagram depicts overlap between all DEG among OTII thymocytes from RIPmOVA v WT hosts and genes induced in CD4 T cells by CA-NFAT-RIT misexpression (PMID: 25680272; filtering as noted in figure). Heatmap depicts expression of overlap genes in our data set. see also Table 2k **e.** Venn diagram depicts overlap between RIPmOVA-induced genes in our data set (white circle, filtering criteria as noted in figure) and CD8 exhaustion signature identified by comparing Tex with anti-PDL1 treated CD8 T cells (PMID: 2778795; filtering as noted in figure). Heatmap depicts expression of overlap genes in our data set and asterisk highlights *Nr4a1/2/3*. see also Table 2l

**Supp Figure 9.**
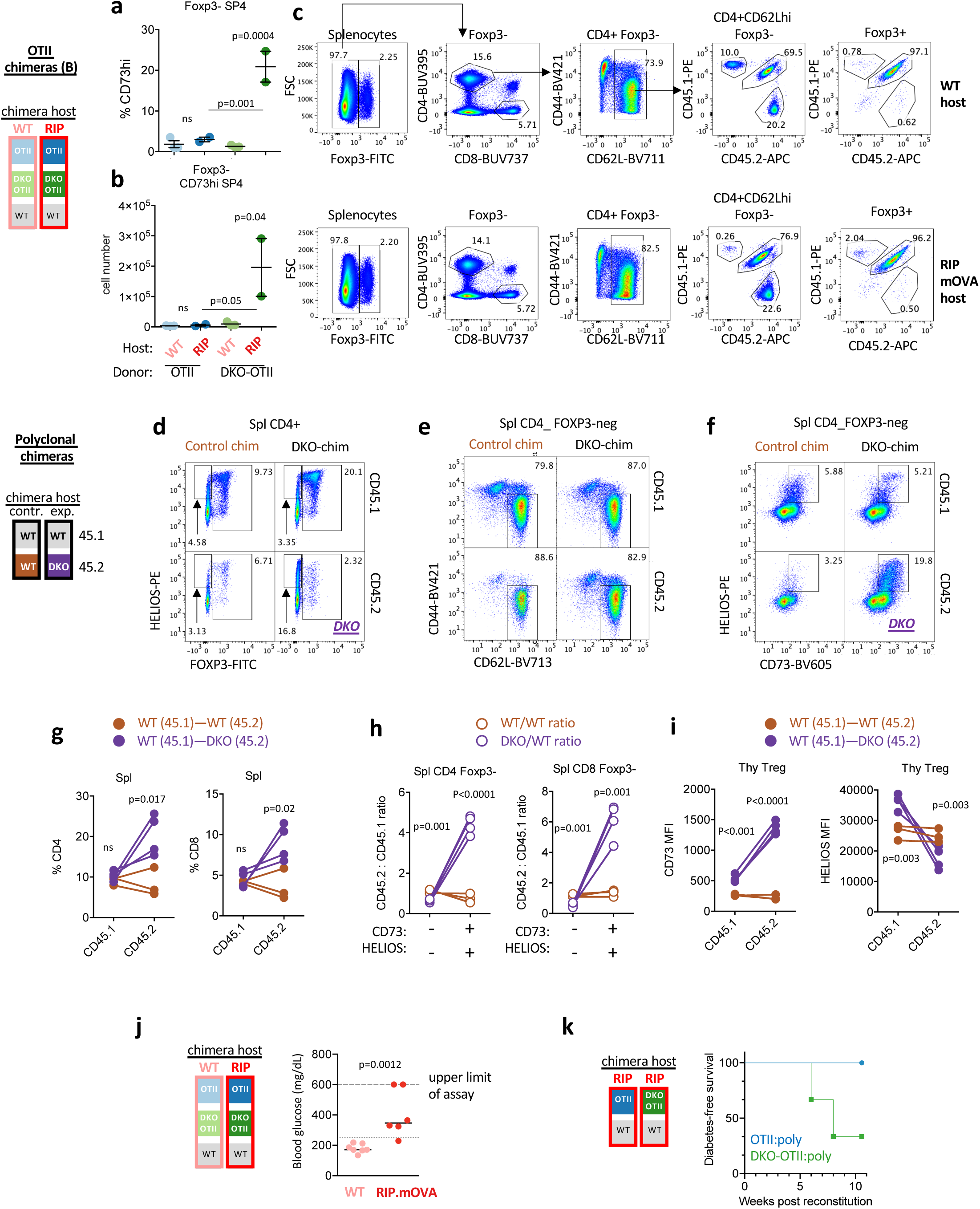
**Related to Main Figures 8, 9** **a, b.** Graphs depict quantification of CD73+ cells among SP4 Foxp3- OTII thymocytes of each genotype from RIPmOVA and WT host chimeras as gated in Fig 8h. Graphs depict N=3 WT host and N=2 RIPmOVA host chimeras +/- SEM. **c.** Gating scheme depicts donor genotype contribution to Foxp3+ CD4 splenocytes and naïve Foxp3- CD4 splenocytes from RIPmOVA and WT host chimeras (see Fig 3a schematic). **d-f.** Representative plots depict splenocytes from polyclonal chimeras (see Fig 1c schematic) stained to identify among each donor genotype intra-cellular FOXP3 and HELIOS within CD4+ gate (d), naïve CD62Lhi CD44lo FOXP3- CD4 T cells (e), and HELIOS+CD73+ gate among FOXP3- CD4 T cells (f). **g.** Graphs depict % CD4 and % CD8 splenic T cells within each donor genotype from polyclonal chimeras (see Fig 1c schematic) as gated in Main Fig 9e. Lines connect donor cells from same chimera. **h.** Graphs depict ratio of donor genotypes in polyclonal chimeras (Fig 1c schematic) from among CD73-HELIOS- and CD73+HELIOS+ FOXP3- splenic T cell populations (as gated in Supp Fig 9f above). **i.** Graphs depict MFI of CD73 (left) and HELIOS (right) in thymic FOXP3+ SP4 Treg from each donor genotype in polyclonal chimeras (Fig 1c schematic). Lines connect donor cells from same chimera. **j.** Blood glucose (mg/dL) from WT or RIPmOVA chimeras (see 3a schematic) at 5.5-6.5 weeks following irradiation and bone marrow transfer. **k.** Survival curve depicts N=3 OTII:poly chimeras in RIPmOVA hosts and N=3 OTII:poly chimeras in RIPmOVA hosts (see 2a schematic). <u>Statistical tests</u>: a, b, one way ANOVA with Tukey’s multiple hypothesis test. Data in c represents N=2 DKO-OTII-RIPmOVA host chimeras and N=3 OTII-RIPmOVA host chimeras and reflect 3 independent sets of chimeras. Data in d-f represent N=4 DKO:WT and N=3 control chimeras. g, h, i unpaired two-tailed t-test, do not assume equal SD j, unpaired t-test, do not assume equal SD (Mann-Whitney)

**Supp Figure 10.**
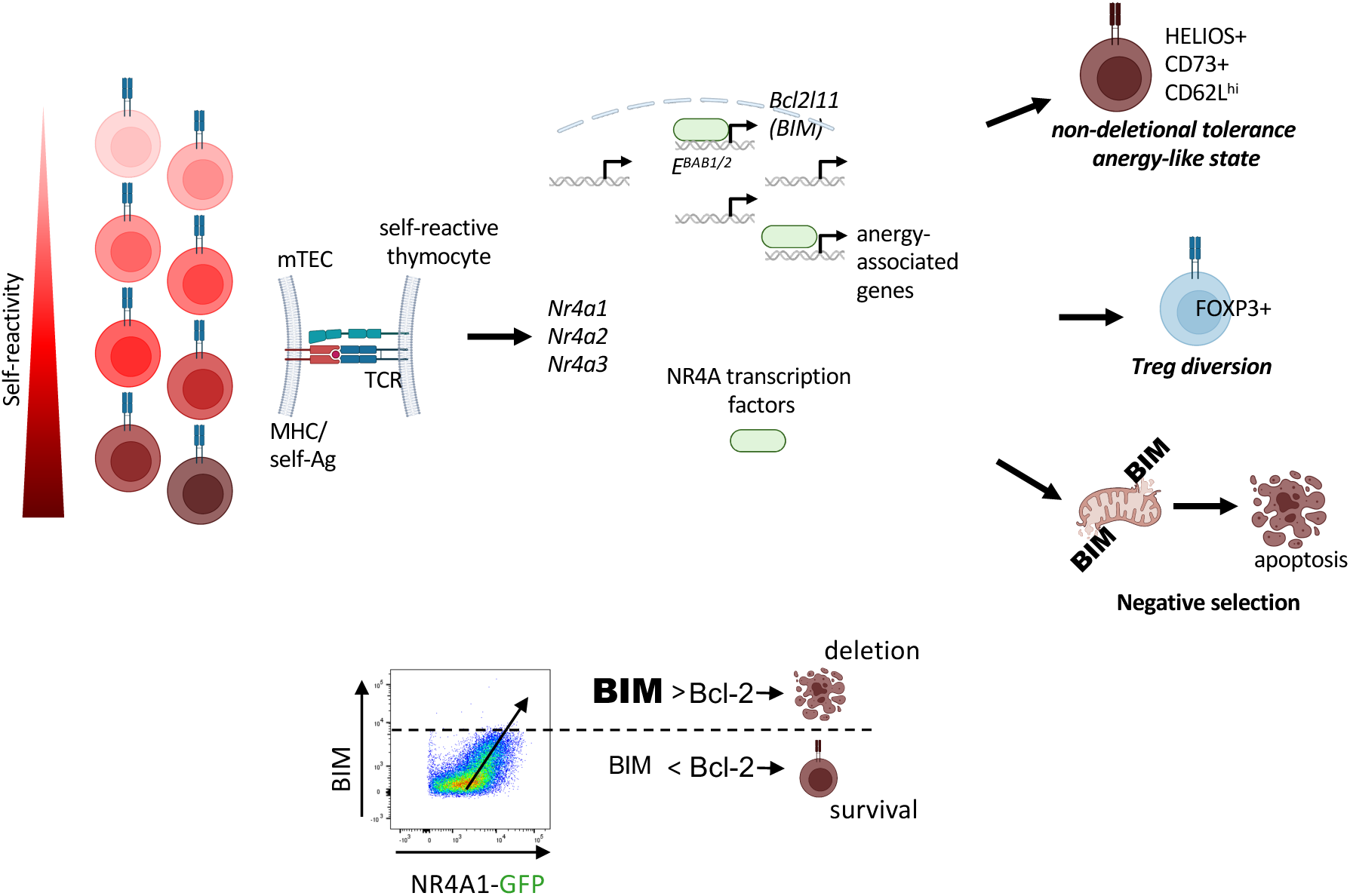
Model. Nr4a genes are TCR-induced in SP thymocytes in the medulla during thymic selection where they induce both *Bcl2l11*/BIM expression via E^BAB^ enhancer engagement and a broader transcriptional program that promotes both Treg diversion and an anergy like program. We hypothesize that when BIM expression stoichiometrically overwhelms anti-apoptotic Bcl2 family members, highly signaled thymocytes with highest Nr4a expression trigger apoptosis. Created with Biorender.com.

## References

1 Husebye, E. S., Anderson, M. S. & Kampe, O. Autoimmune Polyendocrine Syndromes. N Engl J Med 378, 1132–1141 (2018).

2 Anderson, M. S. et al. Projection of an immunological self shadow within the thymus by the aire protein. Science 298, 1395–1401 (2002).

3 Sakaguchi, S. Naturally arising Foxp3-expressing CD25+CD4+ regulatory T cells in immunological tolerance to self and non-self. Nat Immunol 6, 345–352 (2005).

4 Klein, L., Kyewski, B., Allen, P. M. & Hogquist, K. A. Positive and negative selection of the T cell repertoire: what thymocytes see (and don’t see). Nat Rev Immunol 14, 377–391 (2014).

5 Klein, L., Robey, E. A. & Hsieh, C. S. Central CD4(+) T cell tolerance: deletion versus regulatory T cell differentiation. Nat Rev Immunol 19, 7–18 (2019).

6 Kyewski, B. & Klein, L. A central role for central tolerance. Annu Rev Immunol 24, 571–606 (2006).

7 Michelson, D. A., Hase, K., Kaisho, T., Benoist, C. & Mathis, D. Thymic epithelial cells co-opt lineage-defining transcription factors to eliminate autoreactive T cells. Cell 185, 2542–2558 e2518 (2022).

8 Derbinski, J., Schulte, A., Kyewski, B. & Klein, L. Promiscuous gene expression in medullary thymic epithelial cells mirrors the peripheral self. Nat Immunol 2, 1032–1039 (2001).

9 Bouillet, P. et al. BH3-only Bcl-2 family member Bim is required for apoptosis of autoreactive thymocytes. Nature 415, 922–926 (2002).

10 Woronicz, J. D., Calnan, B., Ngo, V. & Winoto, A. Requirement for the orphan steroid receptor Nur77 in apoptosis of T-cell hybridomas. Nature 367, 277–281 (1994).

11 Woronicz, J. D. et al. Regulation of the Nur77 orphan steroid receptor in activation-induced apoptosis. Mol Cell Biol 15, 6364–6376 (1995).

12 Liu, Z. G., Smith, S. W., McLaughlin, K. A., Schwartz, L. M. & Osborne, B. A. Apoptotic signals delivered through the T-cell receptor of a T-cell hybrid require the immediate-early gene nur77. Nature 367, 281–284 (1994).

13 Zhou, T. et al. Inhibition of Nur77/Nurr1 leads to inefficient clonal deletion of self-reactive T cells. J Exp Med 183, 1879–1892 (1996).

14 Cheng, L. E., Chan, F. K., Cado, D. & Winoto, A. Functional redundancy of the Nur77 and Nor-1 orphan steroid receptors in T-cell apoptosis. EMBO J 16, 1865–1875 (1997).

15 Lee, S. L. et al. Unimpaired thymic and peripheral T cell death in mice lacking the nuclear receptor NGFI-B (Nur77). Science 269, 532–535 (1995).

16 Fassett, M. S., Jiang, W., D’Alise, A. M., Mathis, D. & Benoist, C. Nuclear receptor Nr4a1 modulates both regulatory T-cell (Treg) differentiation and clonal deletion. Proc Natl Acad Sci U S A 109, 3891–3896 (2012).

17 Boulet, S. et al. NR4A3 Mediates Thymic Negative Selection. J Immunol 207, 1055–1064 (2021).

18 Hu, Q. N. & Baldwin, T. A. Differential roles for Bim and Nur77 in thymocyte clonal deletion induced by ubiquitous self-antigen. J Immunol 194, 2643–2653 (2015).

19 Hiwa, R., Nielsen, H. V., Mueller, J. L., Mandla, R. & Zikherman, J. NR4A family members regulate T cell tolerance to preserve immune homeostasis and suppress autoimmunity. JCI Insight 6 (2021).

20 Sekiya, T. et al. Nr4a receptors are essential for thymic regulatory T cell development and immune homeostasis. Nat Immunol 14, 230–237 (2013).

21 Sekiya, T. et al. Suppression of Th2 and Tfh immune reactions by Nr4a receptors in mature T reg cells. J Exp Med 212, 1623–1640 (2015).

22 Mullican, S. E. et al. Abrogation of nuclear receptors Nr4a3 and Nr4a1 leads to development of acute myeloid leukemia. Nature medicine 13, 730–735 (2007).

23 Moran, A. E. et al. T cell receptor signal strength in Treg and iNKT cell development demonstrated by a novel fluorescent reporter mouse. J Exp Med 208, 1279–1289 (2011).

24 Zikherman, J., Parameswaran, R. & Weiss, A. Endogenous antigen tunes the responsiveness of naive B cells but not T cells. Nature 489, 160–164 (2012).

25 Zhang, L., Xie, F., Zhang, J., Dijke, P. T. & Zhou, F. SUMO-triggered ubiquitination of NR4A1 controls macrophage cell death. Cell Death Differ 24, 1530–1539 (2017).

26 Huang, B., Pei, H. Z., Chang, H. W. & Baek, S. H. The E3 ubiquitin ligase Trim13 regulates Nur77 stability via casein kinase 2alpha. Sci Rep 8, 13895 (2018).

27 Pei, H. Z., Huang, B., Chang, H. W. & Baek, S. H. Ovarian tumor domain-containing ubiquitin aldehyde binding protein 1 inhibits inflammation by regulating Nur77 stability. Cell Signal 59, 85–95 (2019).

28 Bending, D. et al. A timer for analyzing temporally dynamic changes in transcription during differentiation in vivo. J Cell Biol 217, 2931–2950 (2018).

29 Elliot, T. A. E., et al. Nur77-Tempo mice reveal T cell steady state antigen recognition. Discov Immunol 1, kyac009 (2022).

30 Au-Yeung, B. B. et al. Quantitative and temporal requirements revealed for Zap70 catalytic activity during T cell development. Nat Immunol 15, 687–694 (2014).

31 Au-Yeung, B. B. et al. IL-2 Modulates the TCR Signaling Threshold for CD8 but Not CD4 T Cell Proliferation on a Single-Cell Level. J Immunol 198, 2445–2456 (2017).

32 Au-Yeung, B. B. et al. A sharp T-cell antigen receptor signaling threshold for T-cell proliferation. Proc Natl Acad Sci U S A 111, E3679–3688 (2014).

33 Kishimoto, H. & Sprent, J. Negative selection in the thymus includes semimature T cells. J Exp Med 185, 263–271 (1997).

34 Baldwin, T. A. & Hogquist, K. A. Transcriptional analysis of clonal deletion in vivo. J Immunol 179, 837–844 (2007).

35 Liston, A. et al. Generalized resistance to thymic deletion in the NOD mouse; a polygenic trait characterized by defective induction of Bim. Immunity 21, 817–830 (2004).

36 Steier, Z. et al. Single-cell multiomic analysis of thymocyte development reveals drivers of CD4(+) T cell and CD8(+) T cell lineage commitment. Nat Immunol 24, 1579–1590 (2023).

37 Daley, S. R., Hu, D. Y. & Goodnow, C. C. Helios marks strongly autoreactive CD4+ T cells in two major waves of thymic deletion distinguished by induction of PD-1 or NF-kappaB. J Exp Med 210, 269–285 (2013).

38 Kwan, J. & Killeen, N. CCR7 directs the migration of thymocytes into the thymic medulla. J Immunol 172, 3999–4007 (2004).

39 Ueno, T. et al. CCR7 signals are essential for cortex-medulla migration of developing thymocytes. J Exp Med 200, 493–505 (2004).

40 Weinreich, M. A. & Hogquist, K. A. Thymic emigration: when and how T cells leave home. J Immunol 181, 2265–2270 (2008).

41 Kurobe, H. et al. CCR7-dependent cortex-to-medulla migration of positively selected thymocytes is essential for establishing central tolerance. Immunity 24, 165–177 (2006).

42 Nitta, T., Nitta, S., Lei, Y., Lipp, M. & Takahama, Y. CCR7-mediated migration of developing thymocytes to the medulla is essential for negative selection to tissue-restricted antigens. Proc Natl Acad Sci U S A 106, 17129–17133 (2009).

43 Sawicka, M. et al. From pre-DP, post-DP, SP4, and SP8 Thymocyte Cell Counts to a Dynamical Model of Cortical and Medullary Selection. Front Immunol 5, 19 (2014).

44 Sinclair, C., Bains, I., Yates, A. J. & Seddon, B. Asymmetric thymocyte death underlies the CD4:CD8 T-cell ratio in the adaptive immune system. Proc Natl Acad Sci U S A 110, E2905–2914 (2013).

45 Hemmers, S. et al. IL-2 production by self-reactive CD4 thymocytes scales regulatory T cell generation in the thymus. J Exp Med 216, 2466–2478 (2019).

46 Hu, D. Y., Wirasinha, R. C., Goodnow, C. C. & Daley, S. R. IL-2 prevents deletion of developing T-regulatory cells in the thymus. Cell Death Differ 24, 1007–1016 (2017).

47 Owen, D. L. et al. Identification of Cellular Sources of IL-2 Needed for Regulatory T Cell Development and Homeostasis. J Immunol 200, 3926–3933 (2018).

48 Agle, K. et al. Bim regulates the survival and suppressive capability of CD8(+) FOXP3(+) regulatory T cells during murine GVHD. Blood 132, 435–447 (2018).

49 Liston, A. & Aloulou, M. A fresh look at a neglected regulatory lineage: CD8(+)Foxp3(+) Regulatory T cells. Immunol Lett 247, 22–26 (2022).

50 Kurts, C. et al. Constitutive class I-restricted exogenous presentation of self antigens in vivo. J Exp Med 184, 923–930 (1996).

51 Anderson, M. S. et al. The cellular mechanism of Aire control of T cell tolerance. Immunity 23, 227–239 (2005).

52 Liston, A., Lesage, S., Wilson, J., Peltonen, L. & Goodnow, C. C. Aire regulates negative selection of organ-specific T cells. Nat Immunol 4, 350–354 (2003).

53 Bautista, J. L. et al. Intraclonal competition limits the fate determination of regulatory T cells in the thymus. Nat Immunol 10, 610–617 (2009).

54 Xing, Y., Wang, X., Jameson, S. C. & Hogquist, K. A. Late stages of T cell maturation in the thymus involve NF-kappaB and tonic type I interferon signaling. Nat Immunol 17, 565–573 (2016).

55 Strasser, A., Puthalakath, H., O’Reilly, L. A. & Bouillet, P. What do we know about the mechanisms of elimination of autoreactive T and B cells and what challenges remain. Immunol Cell Biol 86, 57–66 (2008).

56 Cante-Barrett, K., Gallo, E. M., Winslow, M. M. & Crabtree, G. R. Thymocyte negative selection is mediated by protein kinase C- and Ca2+-dependent transcriptional induction of bim [corrected]. J Immunol 176, 2299–2306 (2006).

57 Suen, A. Y. & Baldwin, T. A. Proapoptotic protein Bim is differentially required during thymic clonal deletion to ubiquitous versus tissue-restricted antigens. Proc Natl Acad Sci U S A 109, 893–898 (2012).

58 Stritesky, G. L. et al. Murine thymic selection quantified using a unique method to capture deleted T cells. Proc Natl Acad Sci U S A 110, 4679–4684 (2013).

59 Schmitz, I., Clayton, L. K. & Reinherz, E. L. Gene expression analysis of thymocyte selection in vivo. Int Immunol 15, 1237–1248 (2003).

60 Liston, A. et al. Impairment of organ-specific T cell negative selection by diabetes susceptibility genes: genomic analysis by mRNA profiling. Genome Biol 8, R12 (2007).

61 Gray, D. H. et al. The BH3-only proteins Bim and Puma cooperate to impose deletional tolerance of organ-specific antigens. Immunity 37, 451–462 (2012).

62 Hojo, M. A. et al. Identification of a genomic enhancer that enforces proper apoptosis induction in thymic negative selection. Nat Commun 10, 2603 (2019).

63 Koenis, D. S. et al. Nuclear Receptor Nur77 Limits the Macrophage Inflammatory Response through Transcriptional Reprogramming of Mitochondrial Metabolism. Cell Rep 24, 2127–2140 e2127 (2018).

64 Sekiya, T. et al. Nr4a Receptors Regulate Development and Death of Labile Treg Precursors to Prevent Generation of Pathogenic Self-Reactive Cells. Cell Rep 24, 1627–1638 e1626 (2018).

65 Rajpal, A. et al. Transcriptional activation of known and novel apoptotic pathways by Nur77 orphan steroid receptor. EMBO J 22, 6526–6536 (2003).

66 Liu, X. et al. Genome-wide analysis identifies NR4A1 as a key mediator of T cell dysfunction. Nature 567, 525–529 (2019).

67 Macian, F. et al. Transcriptional mechanisms underlying lymphocyte tolerance. Cell 109, 719–731 (2002).

68 Martinez, G. J. et al. The transcription factor NFAT promotes exhaustion of activated CD8(+) T cells. Immunity 42, 265–278 (2015).

69 Pauken, K. E. et al. Epigenetic stability of exhausted T cells limits durability of reinvigoration by PD-1 blockade. Science 354, 1160–1165 (2016).

70 Kalekar, L. A. et al. CD4(+) T cell anergy prevents autoimmunity and generates regulatory T cell precursors. Nat Immunol 17, 304–314 (2016).

71 Zinzow-Kramer, W. M. et al. Strong Basal/Tonic TCR Signals Are Associated with Negative Regulation of Naive CD4(+) T Cells. Immunohorizons 6, 671–683 (2022).

72 Zinzow-Kramer, W. M., Weiss, A. & Au-Yeung, B. B. Adaptation by naive CD4(+) T cells to self-antigen-dependent TCR signaling induces functional heterogeneity and tolerance. Proc Natl Acad Sci U S A 116, 15160–15169 (2019).

73 Martin, B. et al. Highly self-reactive naive CD4 T cells are prone to differentiate into regulatory T cells. Nat Commun 4, 2209 (2013).

74 Boursalian, T. E., Golob, J., Soper, D. M., Cooper, C. J. & Fink, P. J. Continued maturation of thymic emigrants in the periphery. Nat Immunol 5, 418–425 (2004).

75 Tan, C. et al. NR4A nuclear receptors restrain B cell responses to antigen when second signals are absent or limiting. Nat Immunol 21, 1267–1279 (2020).

76 Melichar, H. J., Ross, J. O., Herzmark, P., Hogquist, K. A. & Robey, E. A. Distinct temporal patterns of T cell receptor signaling during positive versus negative selection in situ. Sci Signal 6, ra92 (2013).

77 Jennings, E. et al. Nr4a1 and Nr4a3 Reporter Mice Are Differentially Sensitive to T Cell Receptor Signal Strength and Duration. Cell Rep 33, 108328 (2020).

78 Lin, B. et al. Conversion of Bcl-2 from protector to killer by interaction with nuclear orphan receptor Nur77/TR3. Cell 116, 527–540 (2004).

79 Thompson, J. & Winoto, A. During negative selection, Nur77 family proteins translocate to mitochondria where they associate with Bcl-2 and expose its proapoptotic BH3 domain. J Exp Med 205, 1029–1036 (2008).

80 Banta, K. L., Wang, X., Das, P. & Winoto, A. B cell lymphoma 2 (Bcl-2) residues essential for Bcl-2’s apoptosis-inducing interaction with Nur77/Nor-1 orphan steroid receptors. J Biol Chem 293, 4724–4734 (2018).

81 Calnan, B. J., Szychowski, S., Chan, F. K., Cado, D. & Winoto, A. A role for the orphan steroid receptor Nur77 in apoptosis accompanying antigen-induced negative selection. Immunity 3, 273–282 (1995).

82 Barron, L. et al. Cutting edge: mechanisms of IL-2-dependent maintenance of functional regulatory T cells. J Immunol 185, 6426–6430 (2010).

83 Chen, J. et al. NR4A transcription factors limit CAR T cell function in solid tumours. Nature 567, 530–534 (2019).

84 Ramsdell, F., Lantz, T. & Fowlkes, B. J. A nondeletional mechanism of thymic self tolerance. Science 246, 1038–1041 (1989).

85 Blackman, M. A. et al. A role for clonal inactivation in T cell tolerance to Mls-1a. Nature 345, 540–542 (1990).

86 Schonrich, G., Momburg, F., Hammerling, G. J. & Arnold, B. Anergy induced by thymic medullary epithelium. Eur J Immunol 22, 1687–1691 (1992).

87 Malhotra, D. et al. Tolerance is established in polyclonal CD4(+) T cells by distinct mechanisms, according to self-peptide expression patterns. Nat Immunol 17, 187–195 (2016).

88 Pobezinsky, L. A. et al. Clonal deletion and the fate of autoreactive thymocytes that survive negative selection. Nat Immunol 13, 569–578 (2012).

89 Ludwinski, M. W. et al. Critical roles of Bim in T cell activation and T cell-mediated autoimmune inflammation in mice. J Clin Invest 119, 1706–1713 (2009).

90 May, J. F. et al. Establishment of CD8+ T Cell Thymic Central Tolerance to Tissue-Restricted Antigen Requires PD-1. J Immunol 212, 271–283 (2024).

91 Fenioux, C. et al. Thymus alterations and susceptibility to immune checkpoint inhibitor myocarditis. Nat Med 29, 3100–3110 (2023).

92 Cunningham, C. A., Helm, E. Y. & Fink, P. J. Reinterpreting recent thymic emigrant function: defective or adaptive? Curr Opin Immunol 51, 1–6 (2018).

93 Barnden, M. J., Allison, J., Heath, W. R. & Carbone, F. R. Defective TCR expression in transgenic mice constructed using cDNA-based alpha- and beta-chain genes under the control of heterologous regulatory elements. Immunol Cell Biol 76, 34–40 (1998).

94 Wan, Y. Y. & Flavell, R. A. Identifying Foxp3-expressing suppressor T cells with a bicistronic reporter. Proc Natl Acad Sci U S A 102, 5126–5131 (2005).

95 Dobin, A. et al. STAR: ultrafast universal RNA-seq aligner. Bioinformatics 29, 15–21 (2013).

96 Lawrence, M. et al. Software for computing and annotating genomic ranges. PLoS Comput Biol 9, e1003118 (2013).

97 Love, M. I., Huber, W. & Anders, S. Moderated estimation of fold change and dispersion for RNA-seq data with DESeq2. Genome Biol 15, 550 (2014).

